# Peptidergic top-down control of metabolic state-dependent behavioral decisions in a conflicting sensory context

**DOI:** 10.1101/2025.03.12.642538

**Authors:** Bibi Nusreen Imambocus, Yushi Jiang, Michael Schleyer, Peter Soba

**Affiliations:** Institute of Physiology and Pathophysiology, Friedrich-Alexander-Universität Erlangen-Nürnberg, 91054, Erlangen, Germany; LIMES Institute, Department of Molecular Brain Physiology and Behavior, University of Bonn, Carl-Troll-Str. 31, 53115, Bonn, Germany; Institute for the Advancement of Higher Education, Hokkaido University, 060-0810, Sapporo, Japan

## Abstract

Animals make economically beneficial behavioral decisions by integrating the external sensory environment with their internal state. The metabolic state, i.e. hunger, can shift priorities, prompting risk-taking and reducing responses to danger during foraging activities. How behavioral changes are processed in the nervous system to generate flexible responses based on needs and risks remains largely unclear. Here, we demonstrate in *Drosophila* larvae that the corticotropin releasing hormone (CRH) homolog diuretic hormone 44 (Dh44) and its producing neurons exert metabolic top-down control of behavioral choice in a conflicting sensory context. The metabolic state via Dh44 signaling mediates the transition from avoiding to tolerating aversive light conditions in the presence of food. In vivo imaging of neuropeptide release revealed that Dh44 regulates the acute release of Insulin-like peptide 7 (Ilp7) specifically in a conflicting context, thereby inducing light tolerance. Optogenetic activation of specific subsets of Dh44 or Ilp7 producing neurons during starvation is sufficient to shift behavior from light tolerance to avoidance by regulating light avoidance circuit responses. This top-down feedforward peptidergic circuit may represent a general mechanism that helps organisms to balance risk-taking with metabolic needs by allowing flexible adjustment of behavior in a conflicting multisensory environment.

## Introduction

All animals are under pressure to continuously update and optimize behavioral decisions. Of particular importance are economic decisions where a valuable entity is weighed against a penalty with the action expected to be beneficial to the animal’s fitness and survival. Economic decisions can vary in magnitude from simple, where the outcome is predictable, to complex, like gambling, involving uncertainty about the benefit of a selected action. Initially proposed by neuroeconomics to be based on absolute values, the concept of decision making has been revisited by neurobiologists ^1^. An optimal evaluation process considers the current sensory context, internal state, learned experiences as well as evolutionary imprints ^2–4^.

Innate behaviors like feeding, sleeping, and escaping from danger are economically valuable for the animal’s physiological state and survival. Feeding is an innate behavior which is particularly important and rewarding to a hungry animal, and it is normally prioritized over other innate behaviors ^5–8^. Hunger is sensed by internal sensors in the body and generates a feeding drive whose purpose is to bring back the system to its homeostatic setpoint.

In mammals, the hypothalamus integrates hunger signals and controls selectivity for innate behaviors through various circuits and neuropeptide release, both in the brain and via humoral responses ^6,9^. Similarly, invertebrates like *Drosophila* possess a neuroendocrine center in the central brain, exerting metabolic control of behavior through local and global signals ^10–13^. Diuretic hormone 44-producing (DH44) neurons are considered to act as internal sensors for nutritive sugars and amino acids promoting food consumption in flies ^14,15^. Diuretic hormone 44 (Dh44) itself is a corticotropin-releasing hormone (CRH) homologue and acts via the G protein coupled receptors Dh44-R1 and Dh44-R2 to regulate feeding responses, rest/activity rhythms, and sleep in flies (Cavanaugh et al. 2014; Dus et al. 2015; Nässel & Zandawala 2022; Poe et al. 2023). It is thus a prime candidate for regulating state-dependent innate behaviors that are prioritized depending on the most urgent physiological need of the organism.

As the feeding drive of animals is under strong selection pressure, metabolic state-dependent modulation of neuronal circuits is primed to affect the behavioral output. Starved zebrafish larvae forage in an otherwise unfavored environment featuring predator-like looming stimuli by dampening the stress response axis and serotonergic recruitment of neurons in the tectum ^18^. Similarly, hungry mice can suppress paw licking due to chronic inflammatory but not acute pain via Neuropeptide Y signaling in the parabrachial nucleus, thus prioritizing the bigger need for feeding in the presence of a food source ^19^. In flies, prioritization for feeding has been shown to dominate sleep drive in starved animals ^20^.

Thus, the coincidence of multiple sensory stimuli with positive and negative valences and competing states need to be integrated by the nervous system to strengthen or dampen specific behaviors. This is particularly true in a conflicting sensory context, where the metabolic state can critically influence behavioral decisions. CO_2_ is aversive for flies, but the presence of food odor inhibits the aversive output pathway in hungry flies ^21,22^. Similarly, conflicting gustatory signals are processed at the sensory or higher order level in a metabolic state-dependent manner, allowing hungry flies to forage on a nutritive food source despite the presence of an aversive bitter substance ^23,24^.

Therefore, goal-oriented acquisition of essential resources like food and potentially competing needs including sleep and shelter has to be computed from the sensory context and the internal state of the animal, with competing innate behaviors allowing for flexible suppression of one another to meet the most urgent need of an animal ^25^. However, it is largely unclear how the feeding drive and internal sensors influence innate circuit dynamics to generate economical decisions in the context of conflicting sensory valences from different modalities.

We designed a behavioral assay that measured the decision of *Drosophila* larvae seeking a risky rewarding or a safe environment depending on their metabolic state. Combining aversive light conditions with food, we found that *Drosophila* larvae chose a safe dark side when sated, but when starved foraged in the risky environment. We showed that DH44 neuron activity and Dh44 signaling exerted top-down control of innate light avoidance in a metabolic state and sensory context-specific manner. We show that Dh44 and the metabolic state control light-induced release of Insulin-like peptide 7 (Ilp7) from dorsal pair Ilp7 (Dp7) neurons, which regulates the light avoidance circuit responses in the presence of nutritive sugar. Activation of downstream neurons was selectively suppressed in the conflicting sensory context in starved animals in a Dp7 neuron-dependent manner. Dampening of the light avoidance enabled starved animals to forage in light, which was reversed in refed animals in a nutrition-dependent manner. Our results thus reveal a mechanism by which the brain adaptively and acutely weighs beneficial and non-beneficial sensory valences in accordance with the metabolic drive to select the most economical behavior.

## Results

### A multisensory choice assay to assess behavior in response to context and state

To test how the metabolic state drives selective behavioral responses in a multisensory context with conflicting stimuli, we designed a suitable choice assay for *Drosophila* larvae (termed multisensory context assay). To this end, we paired the appetitive cue fructose, which has a positive valence and is a preferred sugar for larvae ^26^, with light, which is aversive for larvae and carries a negative valence ^27–30^. We then asked if the paired appetitive and aversive cues fructose and light alter larval preference behavior depending on their metabolic state. Given the choice between the aversive/appetitive pairing and no stimulus (darkness), fed larvae strongly preferred darkness (Fig. 1A). However, animals starved overnight showed a clear preference for fructose despite the aversive light (Fig. 1A). To test if this effect was specific for fructose, we also paired light with glucose as another appetitive sugar. Similarly to fructose, pairing of light with glucose resulted in avoidance in fed animals but was preferred in starved animals (Fig. 1B). We next tested if the nutritional value of the sugar plays a role and paired light with the non-nutritive sugar D-arabinose, which induces preference behavior and supports learning but not survival ^31,32^. In contrast to the nutritive sugars fructose and glucose, fed and starved animals preferred darkness over light and D-arabinose (Fig. 1C). We next tested whether larval behavior is altered in a unisensory context. For fructose only, the metabolic state of the animals did not influence their preference (Fig. 1D), and glucose and arabinose were equally attractive to larvae (Fig. 1E). As previously shown, light alone was aversive in choice assays ^27,28,30^ and resulted in a preferential distribution of animals to the dark side irrespective of their metabolic state (Fig. 1F).

**Figure 1:**
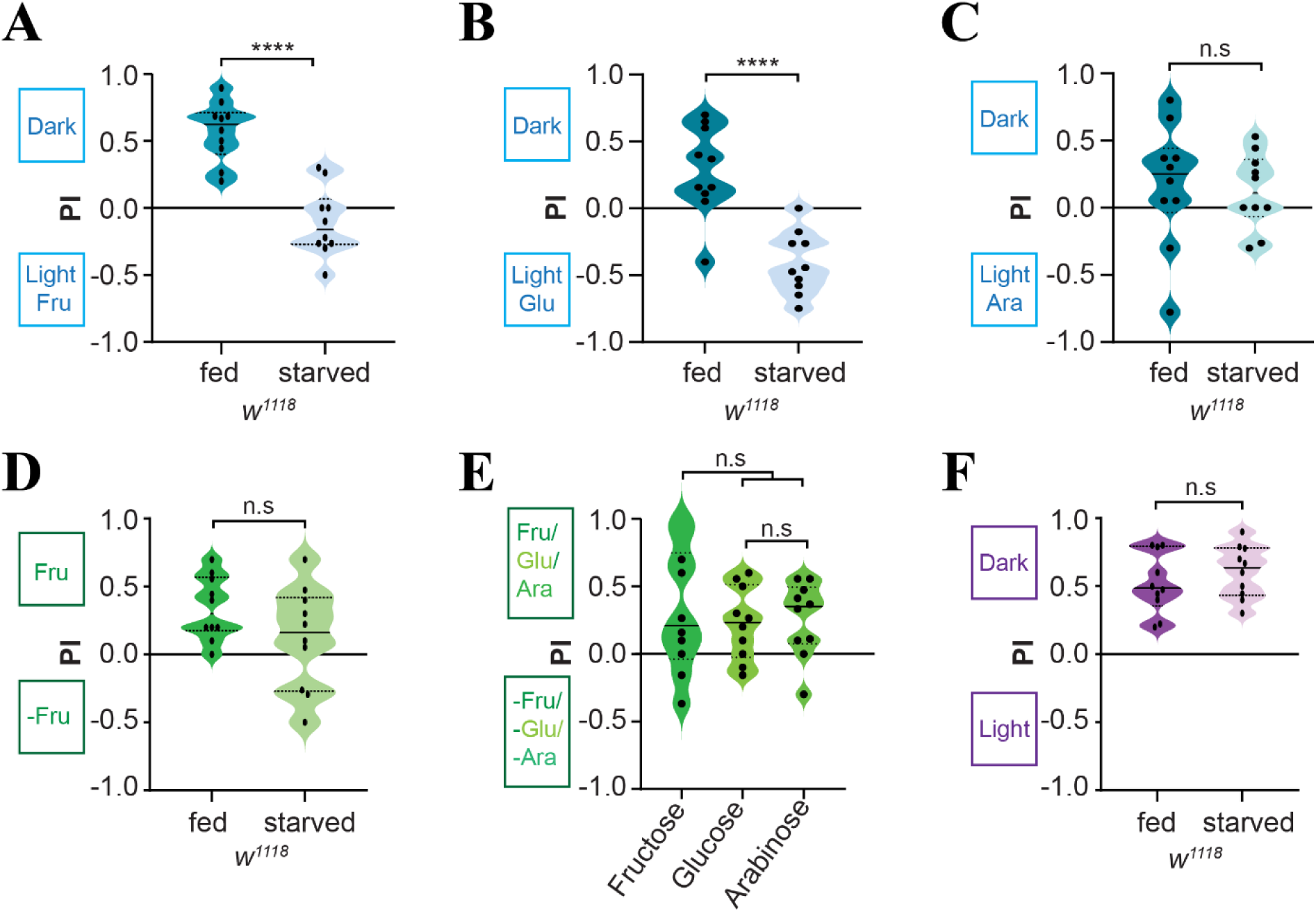
The metabolic state determines behavioral preference in a conflicting sensory context. **A.** Fed larvae prefer darkness while starved larvae show preference for fructose (Fru) in light (n=10 trials, unpaired t-test with Welch’s correction, *** p<0.001). **B.** Fed larvae prefer darkness while starved larvae show preference for glucose (Glu) in light (n= 10 trials, unpaired t-test with Welch’s correction, *** p<0.001). **C.** Fed and starved larvae similarly prefer darkness over arabinose (Ara) in light (n= 10 trials, unpaired t-test with Welch’s correction, n.s. non-significant). **D.** Fed and starved larvae show comparable preference to fructose (Fru) in unisensory preference assays (n=10 trials, unpaired t-test with Welch’s correction n.s. non-significant). **E.** Preference for different sugars (fructose, glucose, arabinose) is similar in fed larvae (n= 10 trials/genotype, one-way ANOVA with Tukey’s posthoc test, n.s. non-significant). **F.** Fed and starved larvae show a similar preference for darkness in unisensory light avoidance assays (n= 10 trials, n.s. non-significant, unpaired t-test with Welch’s correction).

Overall, in addition to the metabolic state, the nutritional value of the appetitive cue in the context of aversive light is relevant for *Drosophila* larvae to show adaptive behavior. Together, our multisensory context assay selectively reveals behavioral adaptation to conflicting sensory stimuli in a state-dependent manner, providing a platform for investigating the neural correlates mediating this behavioral choice.

### DH44 neurons and Dh44 are required for state-dependent behavior

We first set out to identify putative internal sensors controlling the metabolic state of an animal, which should be a key driver of adaptive changes in metabolic state-dependent behavior. The CRH homolog Dh44 has been suggested as a state sensor that regulates feeding and sleep in *Drosophila* ^14,16,33^. In adult flies, independent of the taste modality, DH44 neurons respond to nutritive sugars and selectively guide flies to choose nutritive D-fructose over non-nutritive D-arabinose in gustatory assays (Dus et al. 2015).

As *Dh44* function has not been tested in respect to metabolic state sensing in larvae before, we addressed its role in our multisensory context assay. We tested Dh44 function by investigating the behavioral preference of *Dh44^KO^* as well as *Dh44* receptor (*Dh44R1^KO^* and *Dh44R2^KO^*) mutant animals in multisensory context assays. In all cases, impairing DH44 signaling resulted in increased preference for fructose despite the presence of light (Fig 2A). However, in unisensory light avoidance and fructose preference assays, larvae lacking the *Dh44* or its receptors showed similar behavior as *controls* (Fig 2 B, C). These findings suggest Dh44 signaling is linked to metabolic state-dependent behavioral preferences to appetitive and nutritional sugars despite detrimental light conditions.

**Figure 2:**
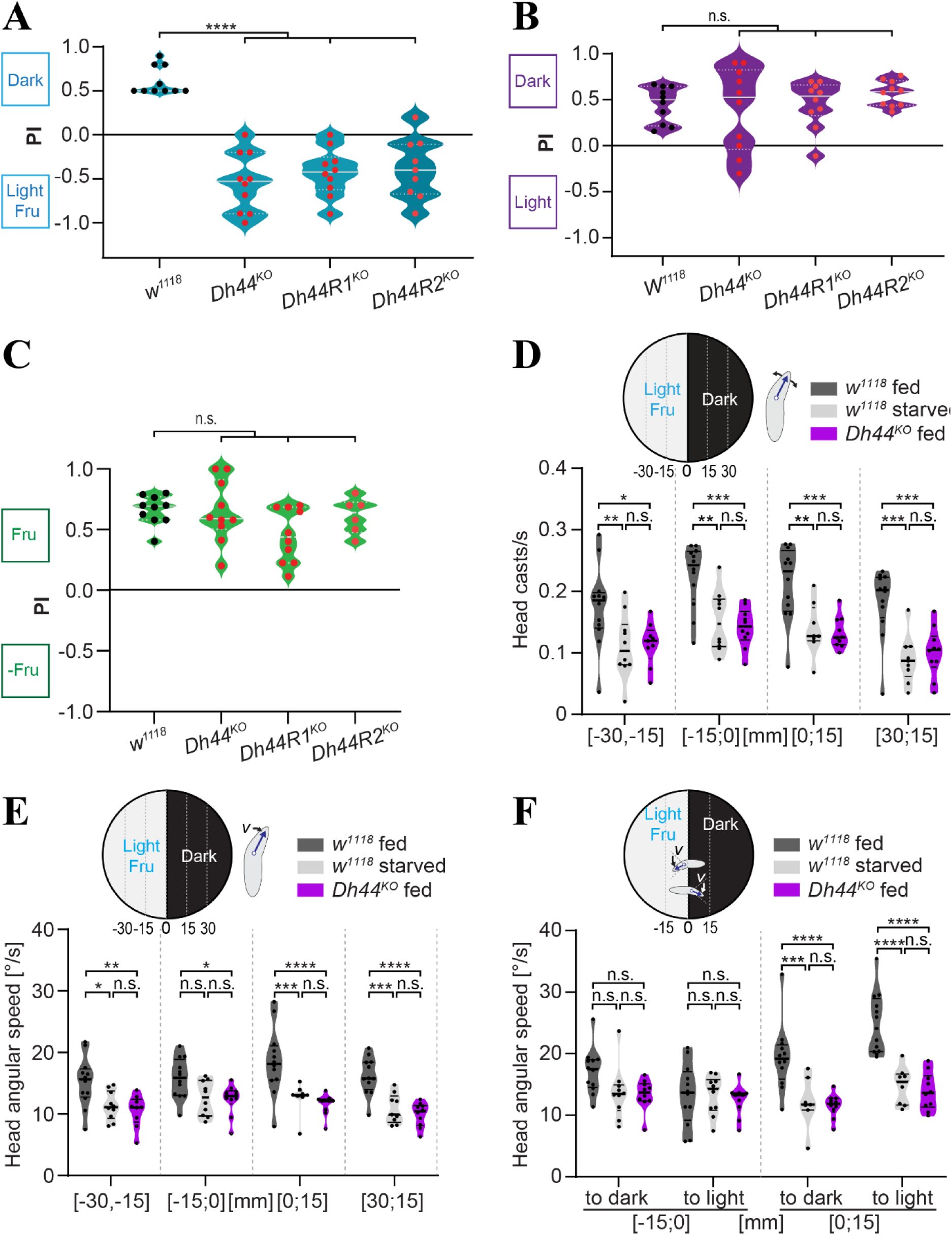
DH44 signaling mediates metabolic state-dependent behavioral switching in a conflicting sensory context. **A.** Fed larvae lacking Dh44 or its receptors Dh44-R1/R2 but not controls prefer foraging on fructose in light (n=10 trials/genotype, one-way ANOVA with Tukey’s posthoc test, ****p<0.0001, n.s. non-significant) **B.** Fed larvae lacking Dh44 or its receptors Dh44R1/R2 display light avoidance behavior comparable to controls (n=10 trials, one-way ANOVA with Tukey’s posthoc test, n.s. non-significant). **C.** Fed larvae lacking Dh44 or its receptors Dh44R1/R2 display fructose preference comparable to controls (n=10 trials, one-way ANOVA with Tukey’s posthoc test, *p<0.05, **p<0.01, ***p<0.001, ****p<0.0001, n.s. non-significant). **D.** Comparison of head casting rate in different zones of the behavioral arena based on automated tracking of larvae during the initial 5min of exploration for fed and starved controls (*w^1118^*) and *DH44^KO^* animals. *DH44^KO^*animals display head casting rates comparable to starved controls, while fed controls display higher head casting rates throughout the arena (n=10 trials, one-way ANOVA with Tukey’s posthoc test, *p<0.05, **p<0.01, ***p<0.001, n.s. non-significant). **E.** Comparison of head angular speed in different zones of the behavioral arena based on automated tracking as in **D**. *DH44^KO^* animals display head angular speed comparable to starved controls, while fed controls display higher values throughout the arena (n=10 trials, one-way ANOVA with Tukey’s posthoc test, *p<0.05, **p<0.01, ***p<0.001, ****p<0.0001, n.s. non-significant). **F.** Comparison of larval head angular speed at the transition zone between dark and light/fructose of animals heading towards or away from the transition zone (to light/dark within 15 mm of the substrate border). Only fed controls but not starved or *DH44^KO^* animals display higher head angular speed on the dark side of the arena (n=10 trials, one-way ANOVA with Tukey’s posthoc test, ***p<0.001, ****p<0.0001, n.s. non-significant).

We further compared behavioral parameters in fed and starved controls and *Dh44^KO^* animals using high-resolution tracking and analyses. We divided the two-choice arena into different segments and analyzed position-dependent larval head casting rate, head angular speed, and midpoint speed (Ext. Fig. 2A-C). While starved larvae were generally slower than fed control animals, they also displayed reduced head casting rates and head angular speed indicating reduced turning behavior. We then binned different regions of the arena relative to the dark/light boundary to delineate position-dependent behavioral differences. Head casting rate and angular speed of fed control animals were increased throughout the arena compared to starved controls (Fig. 2 D,E). Strikingly, fed *Dh44^KO^* animals displayed a locomotion speed comparable to fed control animals (Ext. Fig. 2D), suggesting that the reduced speed of starved larvae was Dh44-independent – possibly due to a lack of energy. In contrast, fed *Dh44^KO^* larvae showed reduced head casting rates comparable to starved control animals (Fig. 2D,E) suggesting that reorientation behavior was Dh44-dependent. We then investigated specific behavioral differences close to the dark/light boundary depending on the position (in the dark or light/fructose) and orientation (towards light or dark) of the animals. Interestingly, fed controls displayed elevated head angular speed specifically on the dark side in proximity of the substrate border, but independent of their orientation (Fig. 2F). In contrast, starved control and *Dh44^KO^* animals did not exhibit increased head angular speed throughout the arena. Larval speed was only mildly position-dependent and generally lower for starved controls only (Ext. Fig. 2D, E). Overall, the in-depth behavioral analysis revealed that in starved and *Dh44^KO^* animals, reorientation behavior at the boundary between dark and light/fructose was reduced compared to fed animals, favoring crossing to the light/fructose side.

### DH44 neuron activity and Ilp7 regulate adaptive behavioral preference

Dh44 is expressed by a small subset of neurons in the *pars intercerebralis* (*pi*), which receive enteric, pharyngeal, and indirect input from various sensory systems ^13,16,34^. We thus asked whether *Dh44* neuron activity is altered by the feeding state of the animals, which would indicate metabolic regulation of their activity and DH44 release. To this end, we used the calcium-dependent nuclear import of *LexA* (CaLexA) system ^35^, which reports calcium-induced activity levels using a transcriptional reporter, in our case GFP. We observed significantly reduced GFP expression driven by the CaLexA system in starved compared to fed animals (Fig. 3A, B). This suggests that DH44 neuron activity is reduced upon starvation, indicating that their activity is metabolically regulated and that DH44 signaling impinges on network function in the larval brain in a feeding state-dependent manner.

**Figure 3:**
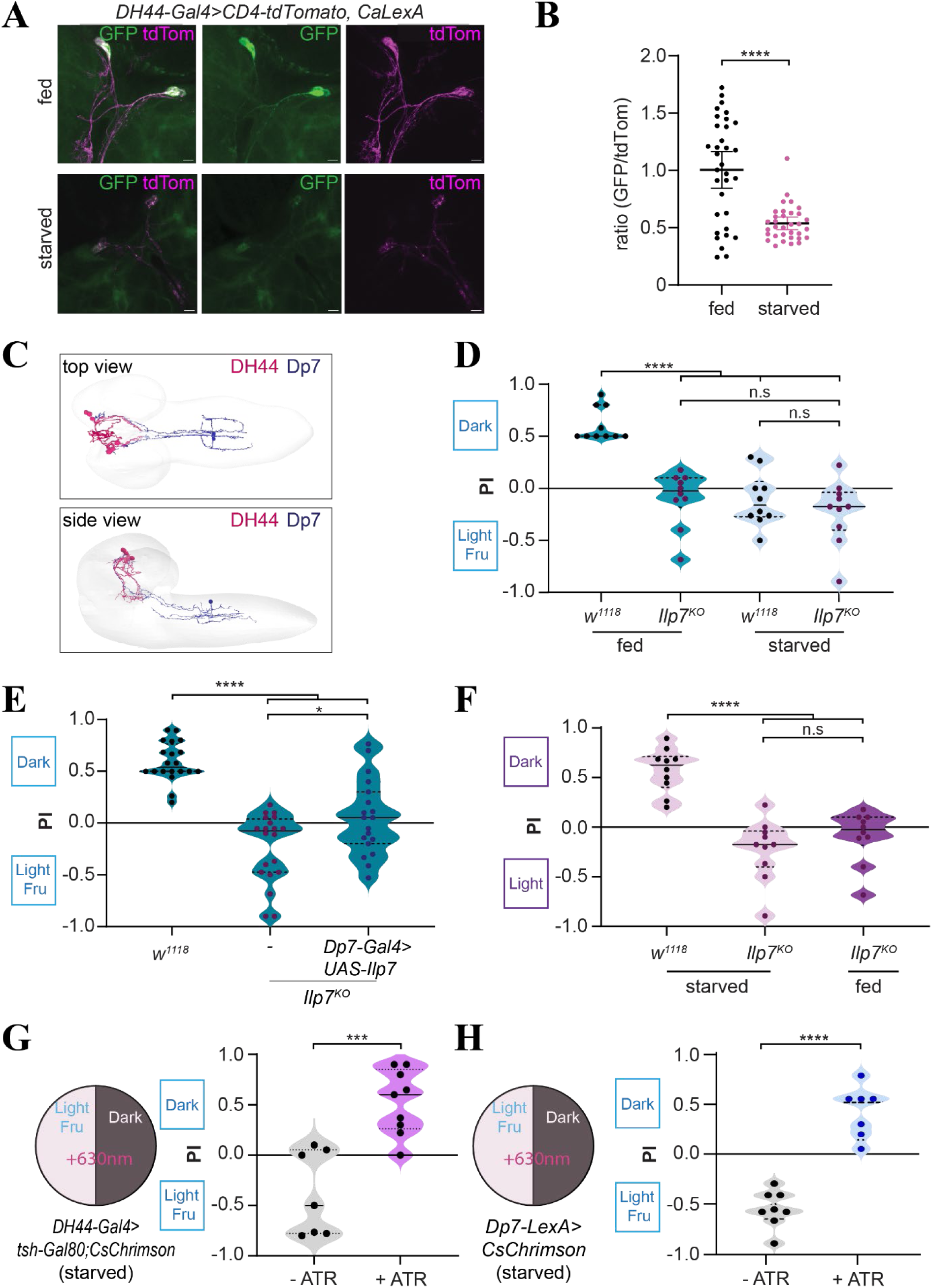
State dependent DH44 neuron activity and Ilp7 signaling regulate behavioral choice in a conflicting sensory context. **A.** Representative images of native CD4-tdTomato and CaLexA-driven GFP signals in DH44 neurons of fed and starved animals (scale bar=10 μm. *Dh44-Gal4>UAS-CD4tdTomato* and *UAS-mLexA-VP16-NFAT, LexAop-GFP*). **B.** Quantification of GFP/tdTomato intensity ratio in DH44 neuron somata in the *pars intercerebralis* (unpaired t-test with Welch’s correction, ****P < 0.0001). **C.** EM reconstruction of Dp7 (blue) and DH44(magenta) neurons (top and side view) showing proximity of arbors in the pi and sez region. **D.** Fed *Ilp7^ko^* but not control larvae show a preference for fructose in light, similar to starved animals of both genotypes (n=10 trials/genotype, one-way ANOVA with Tukey’s posthoc test, ****P < 0.0001, n.s, non-significant). **E.** Ilp7 expression in Dp7 neurons of *Ilp7^ko^* larvae partially restores dark preference in the presence of fructose (n=20 or 10 trials/genotype, one-way ANOVA with Tukey’s posthoc test, ****P < 0.0001, ***P < 0.001, *P < 0.05). **F.** Starved and fed *Ilp7 ^ko^* larvae do not show avoidance in a unisensory light context. (n=10 trials/genotype, one-way ANOVA with Tukey’s posthoc test, ****P < 0.0001, n.s, non-significant). **G.** Optogenetic activation of DH44 neurons in starved animals with and without prior all-trans-retinal feeding (±ATR, *DH44-Gal4, tsh-Gal80, UAS-CsChrimson*, n=7,9, trials, unpaired t-test with Welch’s correction, ****P < 0.0001). **H.** Optogenetic activation of Dp7 neurons in starved animals with and without prior all-trans-retinal feeding (±ATR, *Dp7-LexA, LexAop-CsChrimson*, n=7, 8 trials, unpaired t-test with Welch’s correction, ****P < 0.0001).

We next asked how starved larvae might prioritize fructose foraging over light aversion due to Dh44 signaling. Light avoidance behavior is regulated by multiple sensory neurons and circuits, including Bolwig’s organ and the noxious light-sensitive somatosensory C4da and v’td2 neurons ^27,28,30,36^. Within the somatosensory circuit, dorsal insulin-like peptide 7 (Dp7) neurons play a pivotal role in peptidergic regulation of distinct somatosensory circuits and escape behaviors ^27,37^. In particular, they regulate light avoidance through the acute release of insulin-like peptide 7 (Ilp7). Besides receiving extensive somatosensory input in the ventral nerve cord (VNC), they are ascending neurons innervating the *pi* region harboring the larval neuroendocrine center including DH44 neurons.

We investigated potential direct connections between Dp7 and DH44 neurons, as the latter extend their arbors from the *pi* to the suboesophageal zone (sez) alongside the ascending Dp7 axonal arbor (Fig. 3C). Although no direct synaptic connectivity between Dp7 and DH44 neurons was evident from the larval connectome data ^27,34,38^, their anatomical proximity and the role of Dp7 neurons and Ilp7 in light avoidance prompted us to ask whether they play a role in metabolic control of our context-dependent choice behavior. Under these conditions, we found that *Ilp7^KO^* animals showed increased preference for fructose and light irrespective of their metabolic state and similar to starved controls (Fig. 3D). Rescuing Ilp7 expression specifically in Dp7 neurons (*Ilp7^KO^; Dp7-Gal4>UAS-Ilp7*) resulted in a partial restoration of dark preference in fed larvae (Fig. 3E) suggesting Ilp7 is necessary in Dp7 neurons in this context. Consistently, silencing of Dp7 neurons (*Dp7-LexA*, *LexAop-Kir2.1)* also resulted in preferred distribution of fed animals to the fructose-light side, similarly as starved control animals (Ext. Fig 3A). Under starved conditions, fructose-light preference was observed irrespective of Dp7 neuron silencing, suggesting their activity is not required under these conditions (Ext. Fig 3B).

Dp7 neuron and Ilp7 function have been previously shown to be a part of the innate light avoidance circuit ^27^. Consistently, fed and starved *Ilp7^ko^* larvae displayed similarly reduced dark preference in unisensory light avoidance assays (Fig. 3F). However, fructose preference was not significantly altered upon Kir2.1-mediated silencing of Dp7 neurons in starved animals or in fed and starved *Ilp7^ko^* animals (Ext. Fig. 3C,D). These data confirm that Ilp7 and Dp7 neuron function is required for innate light avoidance but not fructose preference, which did not depend on the animals’ metabolic state under these unisensory conditions.

Due to the confounding role of Dp7 neurons and Ilp7 in innate light avoidance we sought to further test if Dp7 and DH44 neurons can mediate the same adaptive behavioral change in our conflicting sensory context. We therefore asked if acute activation of DH44 or Dp7 neurons is sufficient to change behavior of starved animals in the context of fructose and light. To this end, we optogenetically activated Dh44 neurons in the *pi* or Dp7 neurons in starved animals in a conflicting context, reasoning that increasing their activity and corresponding Ilp7 release might overcome the metabolic state-induced changes. Indeed, under these conditions, CsChrimson-mediated activation of DH44 or Dp7 neurons with red light was sufficient to restore avoidance of the conflicting stimuli in starved larvae (Fig. 3G,H). Together, these data indicate that DH44 and Dp7 neuron activity as well as Dp7-derived Ilp7 function are required to express metabolic state-dependent behavioral adaptation in a conflicting sensory context.

### Context and state render Ilp7 release Dh44-dependent

The findings above suggested that Dh44 and Dp7 neuron activity might be regulated by the metabolic state and sensory context and thus affect Ilp7 release. We thus aimed to investigate Ilp7 release in our conflicting sensory context using our previously characterized Ilp7 neuropeptide release reporter (*NPRR^Ilp7^*), which utilizes an Ilp7 fusion to GCaMP6s to visualize calcium entry upon dense core vesicle fusion with the plasma membrane ^27^. We expressed NPRR^Ilp7^ in Dp7 neurons, which was localized similarly to endogenous Ilp7 and enriched along the axonal arbor within the sez and *pi* region as previously described (Fig. 4A). As Ilp7 can be acutely released from Dp7 neurons upon UV/blue light stimulation ^27^, we imaged intact larvae to monitor Ilp7 release in the *pi* and the sez region of Dp7 neurons, the primary sites of Ilp7 in close proximity to Dh44 neuron arbors (Fig. 4A). To mimic our conflicting context, we imaged UV light-induced Ilp7 release in fed and starved animals in the presence or absence of fructose (Fig. 4B). Interestingly, we detected 3-fold higher Ilp7 peptide release in fed compared to starved animals upon UV light stimulation in the *pi* region (Fig 4C, D). Similarly, UV light-induced Ilp7 peptide release in the sez region was also higher in fed compared to starved animals in the presence of fructose (Ext. Fig 4A, B). In contrast, starved and fed animals subjected only to UV light showed comparable Ilp7 release at both regions (Fig. 4 E, F, Ext. Fig. 4C, D).

**Figure 4:**
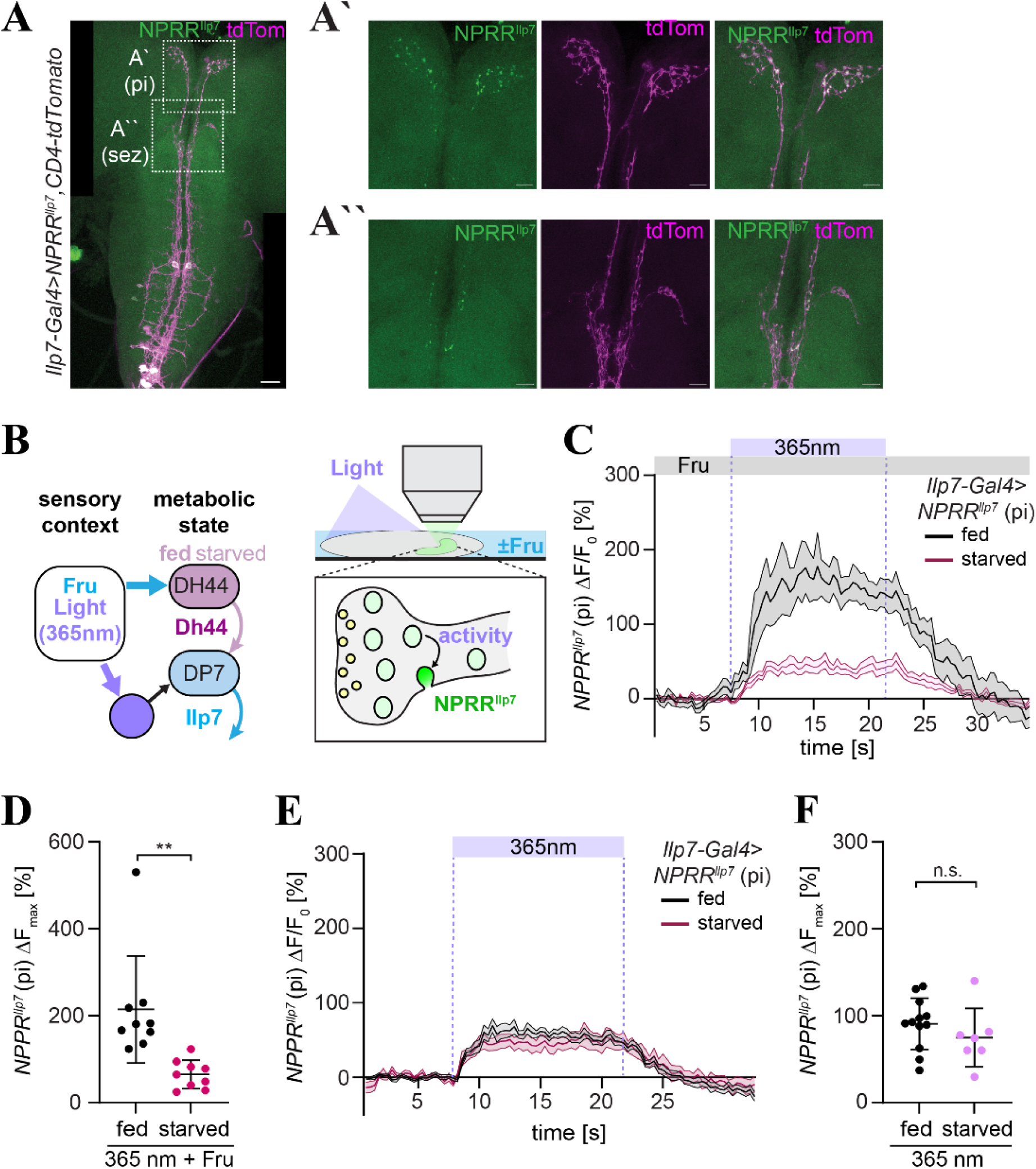
Sensory context and state regulate local Ilp7 release. **A.** Confocal image of 3^rd^ instar larval brain expressing the Ilp7 release reporter (NPRR^Ilp7^) and CD4-tdTomato in Ilp7 neurons (*Ilp7-Gal4, UAS-NPRR^Ilp7^,UAS-CD4tdTomato*). Scale bar=50mm. **À.** Magnified view of NPRR^Ilp7^ and CD4-tdTomato expression in the Dp7 axon terminals in the *pars intercerebralis* (pi). Scale bar=50μm. **À’.** Magnified view of NPRR^Ilp7^ and CD4-tdTomato expression in the Dp7 sez region. Scale bar=50μm. **B.** Schematic of the sensory context (fructose and 365nm light) acting on the metabolic sensor (DH44 neurons) and sensory neurons detecting UV light and needed for Dp7-mediated Ilp7 release. Light was given to intact larvae mounted for confocal imaging of Ilp7 release with or without fructose (±fru). NPRR^Ilp7^ (Ilp7 fused to GCaMP6s) reports Ilp7 release via fluorescence increase after activity-induced fusion of corresponding vesicles with the neuronal plasma membrane. **C.** Average fluorescent change of NPRR^Ilp7^ signals in Dp7 neurons in the pi region (*Ilp7-Gal4>NPRR^ilp7^*) in response to light and in the presence of fructose (0.1M). Fed and starved animals differ in their capacity for Ilp7 release. Dotted lines show the period of stimulation with light (mean±SEM, n=9 neuropeptide puncta from 4 animals). **D.** Maximum fluorescence changes (ΔF_max_) of NPRR^Ilp7^ signals in response to light and fructose in fed and starved larvae (n=9 neuropeptide puncta from 4 animals/genotype, unpaired t-test with Welch’s correction, **P<0.01). **E.** Average fluorescent change of NPRR^Ilp7^ signals in Dp7 neurons in the pi region (*Ilp7-Gal4>NPRR^ilp7^*) in response to light without fructose. Fed and starved animals show similar capacities for Ilp7 release without fructose. Dotted lines show the stimulation period with light (mean±SEM, n=12, 7 neuropeptide puncta from 6 fed and 3 starved animals, respectively). **F.** Maximum fluorescence changes (ΔF_max_) of NPRR^Ilp7^ signals in response to light without fructose in fed and starved larvae (n=12, 7 neuropeptide puncta from 6 fed and 3 starved animals, respectively, unpaired t-test with Welch’s correction, n.s. P>0.05).

These data thus suggest that Ilp7 release is regulated in a metabolic state-dependent manner and only in the presence of the conflicting sensory stimuli (Fig. 5A). Direct comparison of Ilp7 release under all conditions revealed that the presence of fructose increased Ilp7 release only in fed animals. In contrast, release was lower in all other conditions (Fig. 5B). Based on the proximity to DH44 neurons in the *pi* and Dh44-dependent regulation of adaptive behavior in our paradigm, we next investigated whether Dh44 itself could influence Ilp7 release. For this purpose, we imaged UV light-induced Ilp7 release in the presence of fructose in fed DH44 knockout larvae (*Dh44^KO^, Ilp7-Gal4>NPRR^ilp7^*). Interestingly, Ilp7 release in the axon terminals was virtually completely abolished in *DH44^KO^* animals (Fig 5C). In contrast, stimulation with only UV light did not affect Ilp7 release in the *DH44^KO^* background compared to controls (Fig 5D), similarly to the fed/starved conditions investigated before (Fig. 5B). When comparing Ilp7 release activity in the presence or absence of fructose, animals stimulated only with light showed intermediate Dh44-independent release, while the presence of fructose made Ilp7 release Dh44-dependent. These findings suggest that Ilp7 release is regulated by Dh44 only in the conflicting sensory context, consistent with its role in multisensory but not unisensory choice behavior.

**Figure 5:**
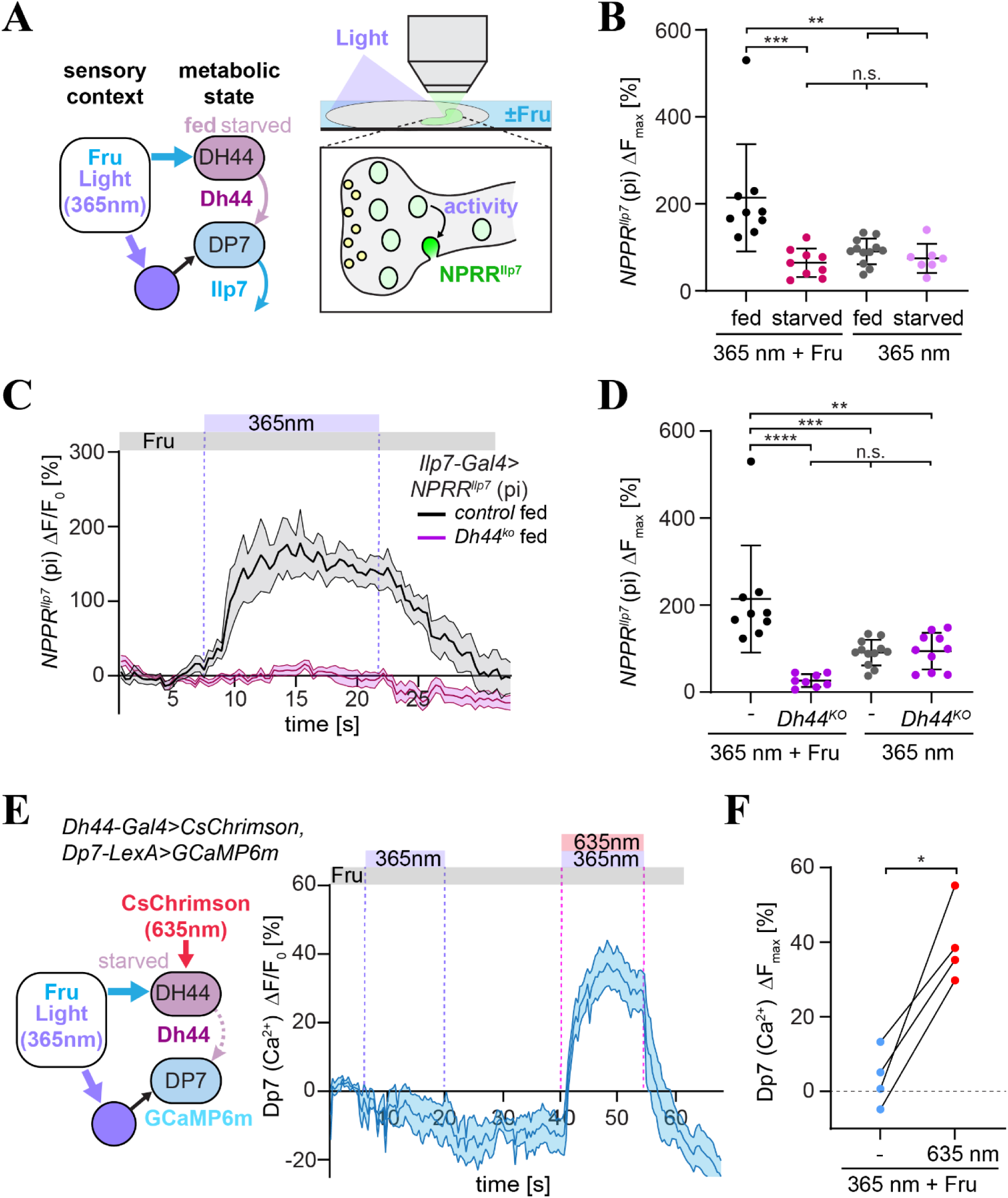
Dh44 is required for Ilp7 peptide release in a context and state-dependent manner. **A.** Schematic of the sensory context (fructose and 365nm light) acting on the metabolic sensor (DH44 neurons) and sensory neurons detecting UV light and needed for Dp7-mediated Ilp7 release. Light was given to intact larvae mounted for confocal imaging of Ilp7 release with or without fructose (±fru). **B.** Comparison of maximum fluorescence changes (ΔF_max_) of NPRR^Ilp7^ signals in response to light in fed and starved larvae with and without fructose (data from Fig. 4 D,F, one-way ANOVA, **P<0.01, ***P<0.001, n.s, non-significant). **C.** Average fluorescent change of NPRR^Ilp7^ signals in Dp7 neurons in the pi region (*Ilp7-Gal4>NPRR^ilp7^*) in response to light and in the presence of fructose (0.1M) in fed control and *Dh44^ko^*animals. Dotted lines show the period of stimulation with light (mean±SEM, n=9,8 neuropeptide puncta from 4 animals/genotype, control data from Fig. 4C). **D.** Maximum fluorescence changes (ΔF_max_) of NPRR^Ilp7^ signals in response to light in fed control and *Dh44^ko^* animals in the presence or absence of fructose (n=9, 8, 12, 10 neuropeptide puncta from 4,4,6,4 a nimals/genotype, one-way ANOVA, **P<0.01, ****p<0.0001, ****p<0.001, n.s, non-significant, control data from Fig. 5B, 4D). **E.** Average fluorescent change of Dp7 neuron calcium (GCaMP6m) signals in starved larvae fed with all-*trans*-retinal in response to 365 nm light and fructose with and without optogenetic activation of DH44 neurons (630 nm, *DH44-Gal4>UAS-CsChrimson; Dp7-LexA,>LexAop-GCaMP6m*, mean±SEM, n=4 animals). **F.** Maximum fluorescence changes (ΔF_max_) of Dp7 calcium signals from E (n=4 animals, paired t-test with Welch’s correction, *p<0.05).

As the metabolic state and lack of Dh44 affected Ilp7 release only in our conflicting context, we wanted to confirm that Dp7 neuron activity and thus Ilp7 release is indeed regulated by Dh44 in starved animals. To this end, we optogenetically activated DH44 neurons and performed calcium imaging of Dp7 neurons in starved animals in the presence of fructose (Fig. 5E). Stimulation with UV light did not elicit significant calcium responses in Dp7 neurons under these conditions, unlike UV light alone as shown previously ^27^. However, simultaneous optogenetic activation of DH44 neurons was sufficient to facilitate Dp7 calcium responses in starved animals in the presence of fructose (Fig. 5E, F). This data confirms that Dh44 and DH44 neuron activity in the *pi* regulate Dp7 neuron activity and Ilp7 release in a context and state-specific manner.

### Metabolic state and context-dependent dampening of light avoidance circuit responses

Previous work showed that abdominal Leucokinin (ABLK) neurons downstream of Dp7 neurons are activated by noxious UV light and concomitant Ilp7 release, which are all required for nociceptive avoidance behavior ^27,39^. As we found that Dp7 neuron activity and Ilp7 release were reduced in starved animals and our conflicting context (see Fig. 5), we wondered if ABLK neuron activity was also altered in a Dp7-dependent manner under these conditions. To this end, we imaged UV light-induced calcium responses in ABLK neurons in starved animals in the presence of fructose while optogenetically manipulating Dp7 neurons (Fig 6A). In the absence of all-*trans*-retinal supplementation, ABLK neurons showed only weak responses, which were, however, strongly increased upon optogenetic Ilp7 neuron activation under the same conditions (Fig. 6B,C). Starvation itself did not influence the responsiveness of ABLK neurons to UV light in the absence of fructose, where they showed comparable calcium responses (Ext. Fig. 6A,B). These findings suggest that ABLK-dependent circuit output depends on state and context-specific Ilp7 neuron activity.

**Figure 6:**
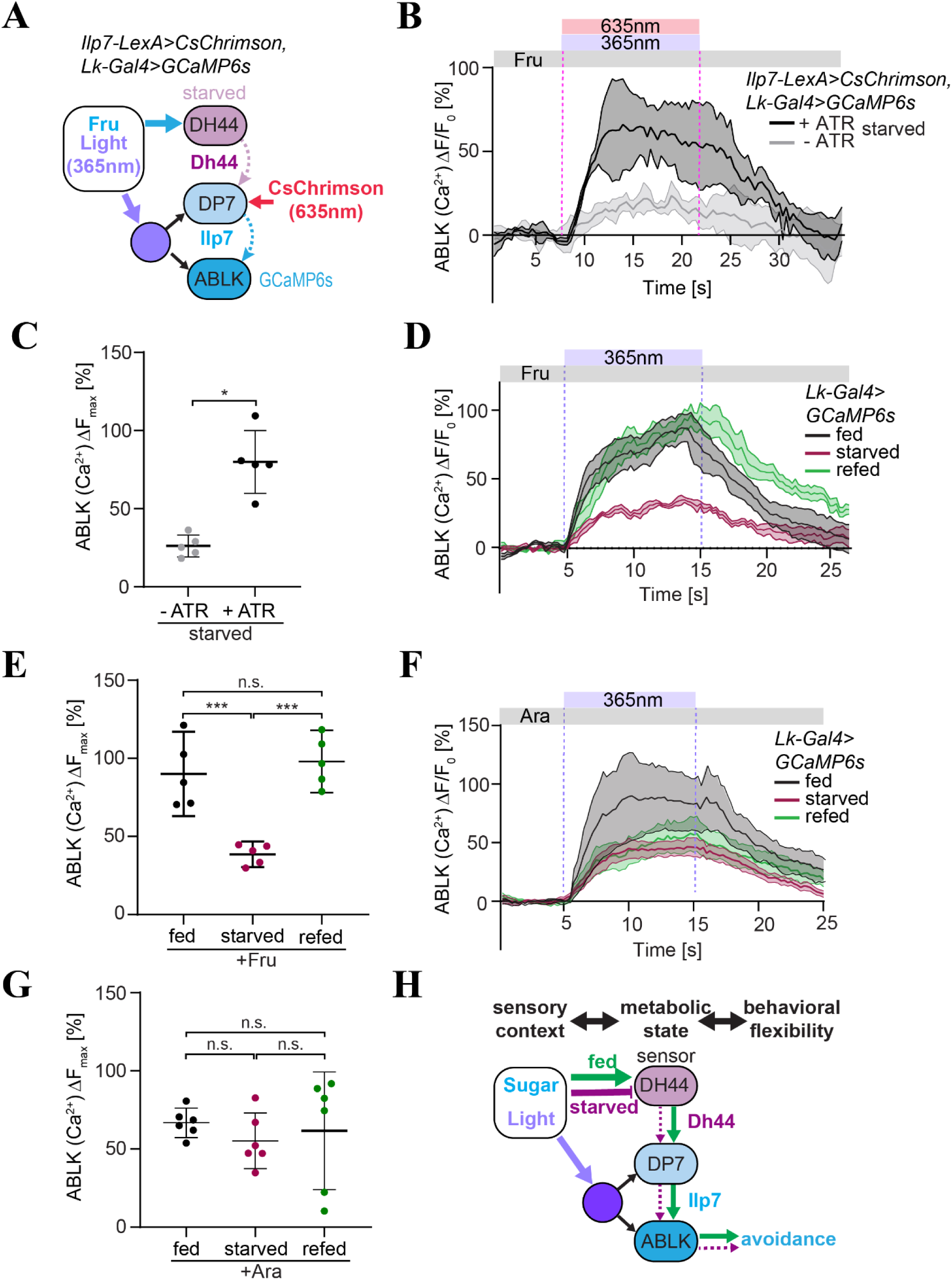
State and context-dependent dampening of light avoidance circuit responses. **A.** Schematic of the sensory context (fructose and 365nm light) acting on the metabolic sensor (DH44 neurons) and sensory neurons detecting UV light acting on Dp7- and downstream ABLK neurons. ABLK neuron calcium responses were imaged in starved larvae with CsChrimson-mediated activation of Dp7 neurons. **B.** Average fluorescent change of ABLK neuron calcium (GCaMP6s) signals in starved larvae in the presence of fructose. Responses to 365 nm light and CsChrimson activation (635 nm) with and without prior all-trans-retinal feeding are shown (±ATR, *Lk-Gal4>UAS-GCaMP6s; Ilp7-LexA>LexAop-CsChrimson*, mean±SEM, n=5 animals). **C.** Maximum fluorescence changes (ΔF_max_) of ABLK calcium signals from **B** (n=5 animals, unpaired t-test with Welch’s correction, *p<0.05). **D.** Average fluorescent change of ABLK neuron calcium (GCaMP6s) signals in fed, starved and refed larvae in response to 365 nm light in the presence of fructose (*Lk-Gal4>UAS-GCaMP6s,* mean±SEM, n=5 animals). **E.** Maximum fluorescence changes (ΔF_max_) of ABLK calcium signals from **D** (n=5 animals, one-way ANOVA, ***p<0.001, n.s, non-significant). **F.** Average fluorescent change of ABLK neuron calcium (GCaMP6s) signals in fed, starved and refed larvae in response to 365 nm light in the presence of arabinose (*Lk-Gal4>UAS-GCaMP6s,* mean±SEM, n=5 animals). **G.** Maximum fluorescence changes (ΔF_max_) of ABLK calcium signals from **F** (n=5 animals, one-way ANOVA, n.s., non-significant). **H.** Model of top-down state-dependent circuit modulation in a conflicting sensory context. Fructose renders Ilp7-dependent noxious light responses sensitive to Dh44. Under starvation noxious light responses in the light avoidance circuit are normal and result in light avoidance. In the presence of the conflicting stimulus fructose, reduced Dh44 action dampens Ilp7 release and light avoidance responses.

To further test if ABLK neuron responses were modulated by the metabolic state in our conflicting sensory context, we imaged UV light-induced calcium responses of ABLK neurons in the presence of fructose in fed and starved larvae. While ABLK responses in fed larvae were comparable to light-only conditions, calcium signals were strongly reduced in starved larvae (Fig. 6D, E, Ext. Fig. 6A, B). We then tested refeeding of starved animals with fructose for 1.5h and observed that ABLK responses to UV light were completely restored in our conflicting sensory context (Fig 6C, D). This finding suggests that the presence of this conflicting context regulates UV light responses of ABLK neurons in a state-dependent manner.

As fructose is a nutritive sugar that also changes the osmolarity of the external solution, which might affect ABLK neuron responses ^39^, we tested ABLK neuron responses to UV light in the presence of D-arabinose, which is non-nutritive and affects osmolarity the same way as fructose. Unlike fructose, the presence of arabinose did not modulate ABLK neuron responses in fed, starved, or arabinose-refed animals (Fig. 6 F, G). This result was consistent with the inability of D-arabinose to induce behavioral changes upon starvation in the conflicting sensory context (see Fig. 1), suggesting that the nutritive value but not osmolarity of the provided sugar is relevant for starvation-induced dampening of ABLK neuron responses.

Taken together, these data show that in a conflicting context of sugar and light, the metabolic state and nutritional value determine the light avoidance circuit responses required for aversive behavior, biasing the behavioral output towards the most urgent need.

## Discussion

Behavioral decisions in a conflicting sensory context have inherent tradeoffs, i. e., the opportunity cost of avoiding foraging in a risky environment by staying safe yet without the chance to feed. While this likely requires value-based decisions, the relative value of two options can vary based on the sensory context, learned valences, and prioritized needs based on the animal’s physiological state ^4,8^. The impact of internal states on driving behavioral outcomes in flies has primarily been studied in individual sensory systems like olfaction and gustation, where hunger can shift odor or taste perception from repulsive to being attractive and vice versa ^21,23,24,40–42^. Our study provides a new platform that integrates multisensory components of different valences to assess adaptive behavior in animals with different internal states. Using light as an aversive stimulus paired with an appetitive sugar, *Drosophila* larvae adapt their behavioral preference according to their metabolic state, which we showed is mediated by Dh44 signaling. Consistently, our in-depth behavioral analysis showed that reorientation behavior, i.e. head casting/angular speed, in starved and *Dh44* deficient animals is altered close to the boundary of the conflicting sensory stimuli of light and fructose. Reorientation is a major mode of larval locomotor behavior to explore and respond to changes in sensory conditions, including light and olfactory stimuli ^43–45^, as well as during associative learning ^46,47^. While fed control larvae displayed elevated reorientation responses close to the fructose/light boundary, consistent with a lower likelihood to cross, starved and *Dh44* deficient animals showed no major change in head casting. Thus, their altered exploratory behavior allows them to freely cross the boundary to forage on the nutritive sugar in light, despite the associated risk of exposure and UV/blue light-induced cellular damage ^48,49^.

As foraging for food is a universal need for survival, growth, and reproduction, many scenarios exist where food availability coincides with increased risk from predators or adverse environmental conditions. In starved flies, feeding overrides courtship drive due to reduced tyramine levels ^50^. Similarly, dopaminergic signaling during *Drosophila* courtship or direct activation of the dopaminergic reward system can overcome a threat or punishment, respectively ^51,52^. While the dopaminergic system might be involved in our case as well, we showed that metabolic state-dependent activity of DH44 neurons and the Dh44 peptide exert top-down control on Ilp7 release and thus the larval light avoidance circuit. Importantly they do so only in a conflicting sensory context and metabolic state-dependent manner.

Internal states are represented by integrated sensory, motor, and internal information that can have multiple physiological and behavioral consequences ^4^. State-inducing neuronal subsets typically integrate different inputs and provide neuromodulatory output acting on many downstream substrates. In *Drosophila*, central neuroendocrine cells including insulin producing cells (IPCs) and DH44 neurons represent such substrates as they can integrate internal and external signals and in turn provide local and humoral responses via their activity level and corresponding peptide release ^14,16,34,53,54^. DH44 neuron activity in the *pi* is regulated by the metabolic state as shown in our work, and they respond to sugar perfusion in starved adult flies ^14,53,55^. While DH44 neurons are presumably involved in postprandial sugar sensing, starved flies display nutritive sugar preference on timescales of less than 1 minute ^14^. DH44 neurons are therefore in a prime position to integrate sensory context and state-dependent signals and exert control over behavior in a top-down manner. A similar function can be attributed to hypothalamic agouti-related peptide (AgRP) neurons in the arcuate nucleus of mammals and connected regions, which are intimately involved in maintaining energy homeostasis and regulation of feeding-related behavior ^9,56^. Activation of AgRP neurons is sufficient to suppress anxiety-like behavior and promote food intake in exposed areas, similarly as observed in starved mice ^57^, and they dampen aggression and fear-related behavior during starvation ^58^. Moreover, the paraventricular nucleus producing CRH and exerting humoral responses via the hypothalamic–pituitary–adrenal (HPA) axis and the lateral hypothalamus in mice respond to the feeding state and are also strongly involved in threat induced physiological responses and behavior ^59,60^. Dh44 exhibits homology to CRH and might similarly not only regulate local circuits but also global physiological responses in a metabolic-state dependent manner ^61–63^.

In this study, we demonstrate that *Drosophila* larvae can context-specifically modulate the salience of aversive responses to align with feeding needs via a top-down control mechanism by the hierarchical action of two critical neuropeptides: Dh44, produced by Dh44-expressing cells in the *pi*, which modulates Ilp7 release from Dp7 neurons only in the presence of nutritive sugar. In turn, Dp7 neuron activity and Ilp7 release are required for contextual Dh44-dependent behavior but also innate light avoidance. Our real-time in vivo approach allowed us to capture the rapid UV light-induced Ilp7 peptide release, acting on short time scales similar to small-molecule neurotransmitters (van den Pol, 2012). While Ilp7 release was unaffected by the metabolic state without a nutritional cue, Dp7 neuron responses and peptide release were rendered Dh44 and feeding state-dependent in our multisensory context. We propose a coincidence detection mechanism ^64,65^, where feeding state-dependent Dh44 action may impact Dp7 neuron activity directly or indirectly, in turn limiting light responses during starvation in a nutritive environment. Consistently, we showed starvation and context-dependent suppression of light avoidance circuit responses in ABLK neurons, which were alleviated by direct optogenetic activation of Dp7 neurons or refeeding of the animals. The activation of ABLK neurons by UV light depends on the synaptic input from a somatosensory circuit and coincidental release of Ilp7, which are essential for light avoidance ^27^. Thus, reduced activity of Dp7 neurons and a corresponding reduction of Ilp7 release under starvation in the presence of a nutritive sugar are fully in line with the observed light avoidance circuit responses. Intriguingly, the non-nutritive sugar D-arabinose, despite inducing preference behavior by itself, was neither able to induce starvation-induced foraging in light, nor modulation of ABLK neuron responses governing light avoidance. This suggests that the ingestion of small sugar amounts is sufficient to detect its nutritional value on a very short time scale, which was previously suggested being dependent on DH44 neurons detecting nutritive sugars and amino acids in adult flies ^14,15^. Consistently, mammalian AgRP neurons are rapidly inhibited within seconds upon sensory detection of food, which further depends on food palatability and the animal’s nutritional state ^66^.

While our results highlight the importance of Dh44-dependent Dp7 activity and Ilp7 release in driving ABLK responses in a context and metabolic state-specific manner, we cannot rule out the potential role of other neuronal components. ABLK and Dp7 neurons also contribute to the scaling and differentiation of different escape behaviors ^27,37,39,67^, which might be affected in the starved state. Nonetheless, our data demonstrate that the delineated peptidergic top-down circuit supports flexible and rapid action selection, allowing animals prioritize behavior in a multisensory context based on feeding needs.

These insights hold significant implications for the neurobiological underpinnings of economic decision-making. Recent evidence suggests that even simple nervous systems, e.g. *C. elegans* with their 302 neurons, can compute economic behavioral decisions when choosing bacterial food based on its nutritional value ^68^. Our data show that *Drosophila* larvae might also make economic decisions by integrating the metabolic state with the nutritional value in a conflicting sensory context, thus optimizing the behavioral outcome to maximize the animal’s benefit. Economic behavior maximizing utility might hence emerge from integrating context and state and subsequent tuning of neuronal circuits through coordinated neuromodulatory effectors to optimize the behavioral outcome.

## Methods

### Fly stock maintenance and rearing

*Drosophila melanogaster* lines and crosses were maintained on standard food at 25°C and 70% humidity with a 12h light /dark cycle. Transgenes were maintained over white mutant (*w^-^*) or yellow-white (*y^-^,w*^-^) backgrounds. Experiments were performed on 3^rd^ instar larvae from males or females aged 94 ± 4 h after egg laying (AEL). Staging of experimental animals was performed as described in ^27^. Briefly, flies were mated in a cage and allowed laying eggs on fresh grape agar plates supplemented with yeast paste for up to 4 hours. Animals were then reared until experimentally used in a humidified petri dish in an incubator at 25°C and 70% humidity with a 12h light /dark cycle. For starvation experiments, 76–80h old larve were subjected to food-deprived conditions. Larvae were transferred to a distilled water-based humidified filter paper in a fly food vial, closed with a vial plug. Starved animals used for optogenetic experiments were continuously kept in the dark and starved by placing 76–80h old larvae on humidified filter paper containing all-*trans*-retinal (5mM).

**Table 1:** Drosophila lines used in this study.

| Reagent or resource | Source | Identifier |
| --- | --- | --- |
| *D. melanogaster: w<sup>1118</sup>* | Bloomington Drosophila Stock Center | BDSC 3605 |
| *D. melanogaster: w<sup>1118</sup>; 20xUAS-GCaMP6s<sup>attP40</sup>* | Bloomington Drosophila Stock Center | BDSC 42746 |
| *D. melanogaster: w<sup>1118</sup>; LexAop-GCaMP6m* | Bloomington Drosophila Stock Center | BDSC 42748 |
| *D. melanogaster: w<sup>1118</sup>; 20XUAS-jGCaMP7s<sup>VK00005</sup>* | Bloomington Drosophila Stock Center | BDSC 79032 |
| *D. melanogaster: w<sup>1118</sup>; 20xUAS-CsChrimson<sup>attP2</sup>* | Bloomington Drosophila Stock Center | BDSC 55136 |
| *D. melanogaster: w<sup>1118</sup>; DH44-R1<sup>KO</sup>* | Bloomington Drosophila Stock Center | BDSC 80700 |
| *D. melanogaster: w<sup>1118</sup>; DH44<sup>KO</sup>* | Bloomington Drosophila Stock Center | BDSC 84492 |
| *D. melanogaster: w<sup>1118</sup>; DH44-R2<sup>KO</sup>* | <sup>69</sup> | N/A |
| *D. melanogaster: w; Ilp7-Gal4* | <sup>70</sup> | N/A |
| *D. melanogaster*: *w*; *Dh44-GAL4* | Bloomington Drosophila Stock Center | BDSC 51987 |
| *D. melanogaster*: *w*; *Calexa* | 71 | N/A |
| *D. melanogaster*: *w*; *Dp7-LexA* | 27 | N/A |
| *D. melanogaster*: *w*; *Ilp7<sup>ko</sup>* | 72 | N/A |
| *D. melanogaster*: *w*; *LexAop-Kir2.1* | 37 | N/A |
| *D. melanogaster*: *w*; *UAS-Kir2.1* | 73 | N/A |
| *D. melanogaster*: *w*; *Dp7-Gal4* | 27 | N/A |
| *D. melanogaster*: *w*; *UAS-NPRR<sup>Ilp7</sup>* | 27 | N/A |
| *D. melanogaster*: *w*; *UAS-Ilp7* | 27 | N/A |
| *D. melanogaster*: *w*; *Lk-Gal4* | 74 | N/A |
| *D. melanogaster*: *w*; *tsh-gal80* | 75 | N/A |

### Conflicting sensory context assay

We established a multisensory conflicting context assay to test whether the feeding state of larvae affects their choices in a need-prioritized manner. The experimental setup was previously used for light avoidance behavior ^27,76^. It consists of a dark chamber with a white light source illuminating one-half of a 10 cm plate while the other half remains dark. The white light source emitted wavelengths ranging from 365 to 580 nm (6.9 to 3.3 μW/mm^2^), while the dark side had less than 0.01 μW/mm^2^ of light.

Divided 10 cm plates (Sarstedt, Nümbrecht, Germany) were used with 2% agar in one, while the other compartment contained 2M fructose, each dissolved in 2% agar in deionized water (C. Roth, Karlsruhe, Germany). Fed or starved larvae aged 94±4h AEL were preincubated in the dark by placing them on a 2% agar plate in a dark chamber for 10 mins. To perform the assay, 17-20 larvae were used per plate. They were placed in the middle of the junction between the light side with fructose and the dark side. Their behavior was recorded for 16 minutes using a Basler ace2040gm camera at 10 frames/s with the Pylon software interface (Basler, Switzerland).

The preference index (PI) was calculated at 15 minutes as PI = (number of larvae in dark - number of larvae in light and fructose)/total number of larvae. PI data was plotted as violin plots, with the central line representing the median. Trials were discarded if more than three larvae escaped from the arena.

Conflicting sensory context assays were also performed using D-glucose (C. Roth, Karlsruhe, Germany)) or D-arabinose (Thermofisher Scientific, Germany). The light-exposed compartment was paired with 2M glucose or 2M arabinose in 2% agar and assays were performed as for fructose above.

### Unisensory light avoidance assay

The experimental chamber used is the same as in the assay described above. Before the experiment larvae were preincubated to dark for 10 mins on a 10 cm 2% agar plate (12 ml of 2% agar dissolved in deionized water, C. Roth, Karlsruh). For each trial, 17-20 animals were placed at the center of the light-dark border and recorded for 16 minutes using a Basler ace2040gm camera at 10 frames/s (Basler, Switzerland). The preference index (PI) was calculated at 15 minutes as PI = (number of larvae in dark - number of larvae in light) / total number of larvae. PI data were plotted as violin plots, with the central line representing the median. Trials were discarded if more than three larvae escaped.

### Unisensory fructose preference assay

Fructose preference assays were performed in the experimental chamber without light illumination using divided plates as above. Animals were assayed for fructose preference in the dark by subjecting them to the center of a 10 cm plate with 2M fructose dissolved in 2% agar on one side versus plain agar (2%) on the other. 17-20 animals were recorded per trial as described above. Preference Index was calculated as above.

### Optogenetic activation behavioral assays

Optogenetic assays were performed on third instar larvae aged 94±4h using CsChrimson-mediated activation of DH44 (*DH44-Gal4*) or Dp7 (*Dp7-LexA*) neurons. For CsChrimson-mediated activation, animals aged up to 24h were supplied with fresh yeast paste containing 5mM all-*trans-*retinal (ATR). After staging, animals were kept at 25^0^C in the dark. Starvation was performed by transferring animals onto a filter paper moistened with either 5mM ATR diluted in ddH_2_O or ddH_2_O only for control animals and incubated o/n in the dark at 25^0^C. Selection of the correct genotype and handling of animals before the assays were done under low red-light illumination. 17-20 larvae were assayed on a divided 10cm petri dish (C. Roth, Karlsruhe, Germany) containing 2% agar/ddH_2_O on one side and 2M fructose dissolved in 2% agar/ddH_2_O on the other side. The experiments for the behavioral assay were performed in the dark chamber as described above, using a white light source to illuminate the fructose side from below. In addition, 635nm light (2-3 μW/mm^2^) was supplied from above on both dark and light side of the arena (CoolLED PE4000). Larvae were cautiously placed in the center of the middle of the petri dish and the behavioral choice of the animals was recorded for 16 minutes. 6-10 trials were performed for each genotype reared under control or experimental conditions. Statistical significance was assessed by t-test with Welch’s correction (GraphPad, San Diego, CA, USA). Violin plots showed the distribution of the animals at 15 mins.

### Animal tracking and behavioral analysis

Recorded videos from the conflicting sensory context assay were tracked and analyzed with IMBA (Individual Maggot Behaviour Analysis) as described in Thane et al.^77^. In brief, the software automatically tracks 12 equidistant spine points from the tail to the head of each larva and can calculate many attributes for each animal. For the current analysis, only the first 5 minutes of each video were included, during which preference is established. At later time points most larvae stayed at the Petri dish border where light reflections prevented reliable tracking. Specifically, we determined the following variables:

- Head angular speed [°/s]: the angular speed of a vector through the head of each animal.
- Head casting rate [head casts/s]: The number of head casting events, lateral head movements associated with turning, per second. A head cast was defined by a head angular speed above a pre-set threshold of 35 °/s, following Thane et al.^77^.
- Midpoint speed [bl/s]: the speed of the midpoint of each larva. As larval speed depends on the size of the animal^77^, we normalized the midpoint speed to the body length (bl) of each individual animal.

Whenever the behavior of larvae is displayed across the position in X direction as line plots, the variable is calculated across all animals of an experimental condition per 1 mm bin, with a sliding averaging filter of 5 mm. For the head casting rate, the head casts per second of each bin are displayed. For the other variables, the mean and the 95 % confidence intervals of the respective variable are displayed.

Whenever data are displayed in violin plots to compare between experimental conditions, we first separated the data in 15 mm wide stripes along the X axis, and calculated the mean of each variable for all animals in each video within this stripe. For some figures, we additionally separated the data according to the direction the animals were heading to. Animals were considered heading “towards dark” whenever the angle between their body orientation and a vector pointing directly towards the dark side (x=1,y=0) was 45° or lower. Animals were considered heading “towards light” when that angle was 135° or higher.

### Larval brain dissection and imaging

Experimental crosses were set up with flies carrying *DH44-Gal4,UAS-CD4tdTomato* and *UAS-CaLexA* (*UAS-mLexA-VP16-NFAT, LexAop-GFP*). Staged larvae aged 94 ± 2h AEL were used for CaLexA experiments. Starvation was performed overnight for 14-16h on dampened filter paper in a test tube which was subsequently incubated at 25°C/70% humidity.

Fed and starved larvae were dissected, and their brains were fixed with 4% PFA in PBS for 15-30 minutes, washed with PBST and mounted on poly-L-lysine coated coverslips (Sigma) in Slow Fade Gold. Native CaLexA-driven GFP and tdTomato signals from DH44 neurons were acquired by confocal microscopy (Zeiss LSM900AS2). The intensity of the CaLexA-driven GFP signal was normalized with the intensity of tdTomato in DH44 neurons. Z-stack images were acquired with the ZEN software (Zeiss) and analyzed using ImageJ (NIH, Bethesda).

Larval brains from genotype *NPRR^ilp7^>Ilp7-Gal4, UAS tdTomato* labeling Ilp7 neuropeptide punctae in Dp7 neurons were dissected in 4% PAF, washed three times with PBST ((PBS with 0.3% Triton X-100 (Roth, Karlsruhe Germany) and mounted on Superfrost slides in Slow Fade Gold (Thermo Fisher, Carlsbad, CA, USA). Native reporter fluorescence of NPRR^ilp7^ was sufficiently bright for visualization under confocal microscopy (Zeiss LSM900). Confocal Z-stacks were processed in Fiji (ImageJ, NIH, Bethesda).

### In vivo calcium imaging

Neuronal activity was recorded live on neuronal soma expressing the calcium sensor GCamP6s/m under the control of specific neuronal Gal4-drivers (*Ilp7-LexA, Lk-Gal4*). Live 3^rd^ instar fed or starved larvae (94±2hrs) were mounted on either 50% glycerol (for unisensory light stimulation) or 100 mM fructose or arabinose in 50% glycerol (for multisensory stimulation) and immobilized with a coverslip. Calcium signals in the cell soma were imaged by confocal microscopy with a 40x/NA1.3 oil objective (Zeiss LSM900AS2). 200 or 400 frame time series were acquired at a frame rate of 0.34s (240 x 240 pixels). Larvae mounted in 50% glycerol/PBS with or without 100 mM fructose or arabinose were illuminated with at least two pulses of UV light for 10s (365-525 nm, 10-60 μW/mm^2^, CoolLED) with an interval of at least 15s between pulses to activate the light avoidance circuit. To restore the feeding state, starved animals were refed for 1.5h with either 1mM fructose or arabinose and imaged.

Optogenetic activation and calcium imaging were performed as described above using CsChrimson-mediated activation of Ilp7 (*Ilp7-LexA*) or Dh44 (*DH44-Gal4*) expressing neurons. CsChrimson was selectively activated with a 630 nm light pulse with or without activation of light avoidance circuit responses (365 nm, 10-60 μW/mm^2^ CoolLED)

Calcium imaging was performed 4-10 larvae/genotype and between ZT 3 to 6. Identical confocal settings consisting of single-plane imaging (approximately 2μm thickness) were used for imaging experiments. Datasets were retained for analysis only if there was no significant focal drift and movement, including a stable baseline and a return to baseline levels after stimulation.

### NPRR^Ilp7^ release imaging

To visualize NPRR^Ilp7^ release, fed or starved third stage larvae (94±2hrs) were mounted in 50% glycerol/PBS with or without100mM fructose for imaging Ilp7 release upon UV light stimulation. Imaging was done as described above. Following a period of baseline imaging for 10 s, light was provided for 10 s (365-525 nm, 10-60 μW/mm^2^ CoolLED) and response decay was imaged for at least 10 s. 2 pulses of light with an interval of at least 15 s were given per given animal. Intensities of individual NPRR^Ilp7^ puncta were quantified over time at the terminal axonal arbors in the *pi* and at the sez region.

### Electron microscopy data set

The morphology and proximity of DH44 and Dp7 neurons were extracted and analyzed from the larval connectome using CATMAID ^78^. The reconstructed DH44 and Dp7 neurons were previously described ^27,34,38^.

### Statistical analysis

Data distribution was tested using a Shapiro-Wilkinson test. If normality was given, a two-tailed unpaired t-test was used to compare 2 groups. For multiple comparisons behavioral assays and in vivo calcium imaging data sets, one-way ANOVA with Tukey’s *post hoc* comparison was performed. Statistical differences were considered significant for *P < 0.05* (*\*P<0.*05, *\*\*P<0.01*, *\*\*\*P<0.001*, *\*\*\*\*P<0.0001*). Statistical testing was performed using GraphPad Prism (GraphPad, San Diego, CA, USA).

Data sets with 2 genotypes were assessed for statistical significance with a t-test with Welch’s correction, and those having more than 3 genotypes were subjected to a one-way ANOVA statistical test with Tukey’s correction (GraphPad, San Diego, CA, USA).

## Acknowledgements

We thank A. Schoofs and M.J. Pankratz for technical advice, reagents and lab space. Stocks obtained from the Bloomington *Drosophila* Stock Center (NIH P40OD018537) were used in this study. This work was supported by grants from the Deutsche Forschungsgemeinschaft (DFG SO 1337/7-1 to PS) and the DFG Heisenberg program (SO1337/6-1 to PS).

## Author contributions

BNI performed and analyzed functional in vivo imaging and behavioral assays, YY performed a subset of behavioral assays, MS performed high resolution tracking and behavioral analysis, BNI and PS designed the study and wrote the manuscript with input from all authors.

## Competing Interests

The authors declare that no competing interests exist.

---

**Extended Figure 2:**
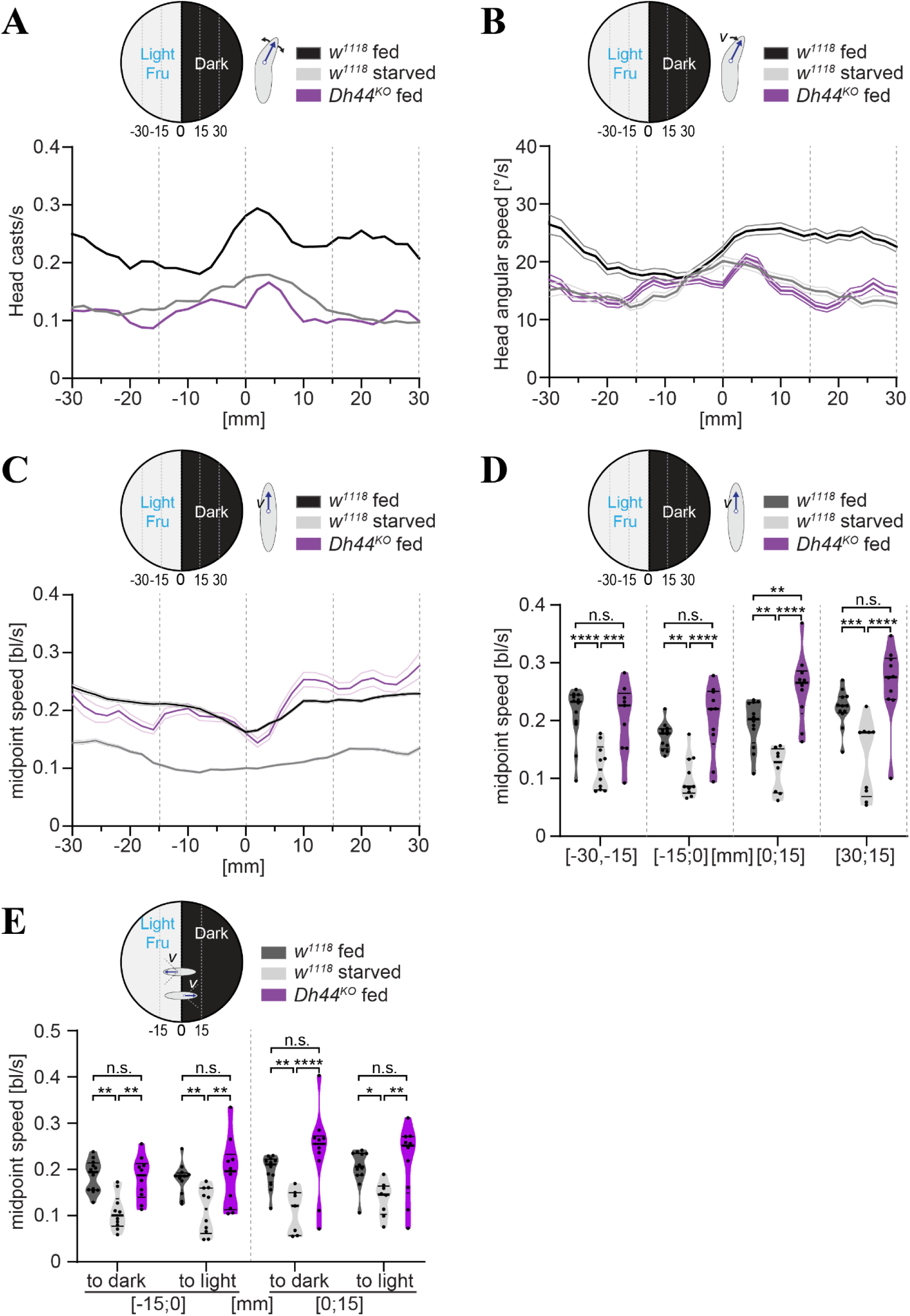
Dh44 signaling mediates metabolic state-dependent behavioral changes in a conflicting sensory context. **A.** Head casting rate in different zones of the behavioral arena based on automated tracking of larvae during the initial 5min of exploration for fed and starved controls (*w^1118^*) and *DH44^KO^* animals. *DH44^KO^* animals display head casting rates comparable to starved controls, while fed controls display higher head casting rates throughout the arena. **B.** Head angular speed in different zones of the behavioral arena based on automated tracking of larvae during the initial 5min of exploration for fed and starved controls (*w^1118^*) and *DH44^KO^* animals. *DH44^KO^*animals display head angular speed comparable to starved controls, while fed controls display higher head angular speed (mean±95% CI). **C.** Midpoint speed in different zones of the behavioral arena based on automated tracking of larvae during the initial 5min of exploration for fed and starved controls (*w^1118^*) and *DH44^KO^* animals (mean±95% CI). **D.** Comparison of midpoint speed in different zones of the behavioral arena based on automated tracking as in **C**. (n=10 trials, one-way ANOVA with Tukey’s posthoc test, **p<0.01, ***p<0.001, ****p<0.0001, n.s. non-significant). **E.** Comparison of larval midpoint speed at the transition zone between dark and light/fructose of animals heading towards or away from the transition zone (n=10 trials, one-way ANOVA with Tukey’s posthoc test, *p<0.05, **p<0.01, ****p<0.0001, n.s. non-significant).

**Extended Figure 3.**
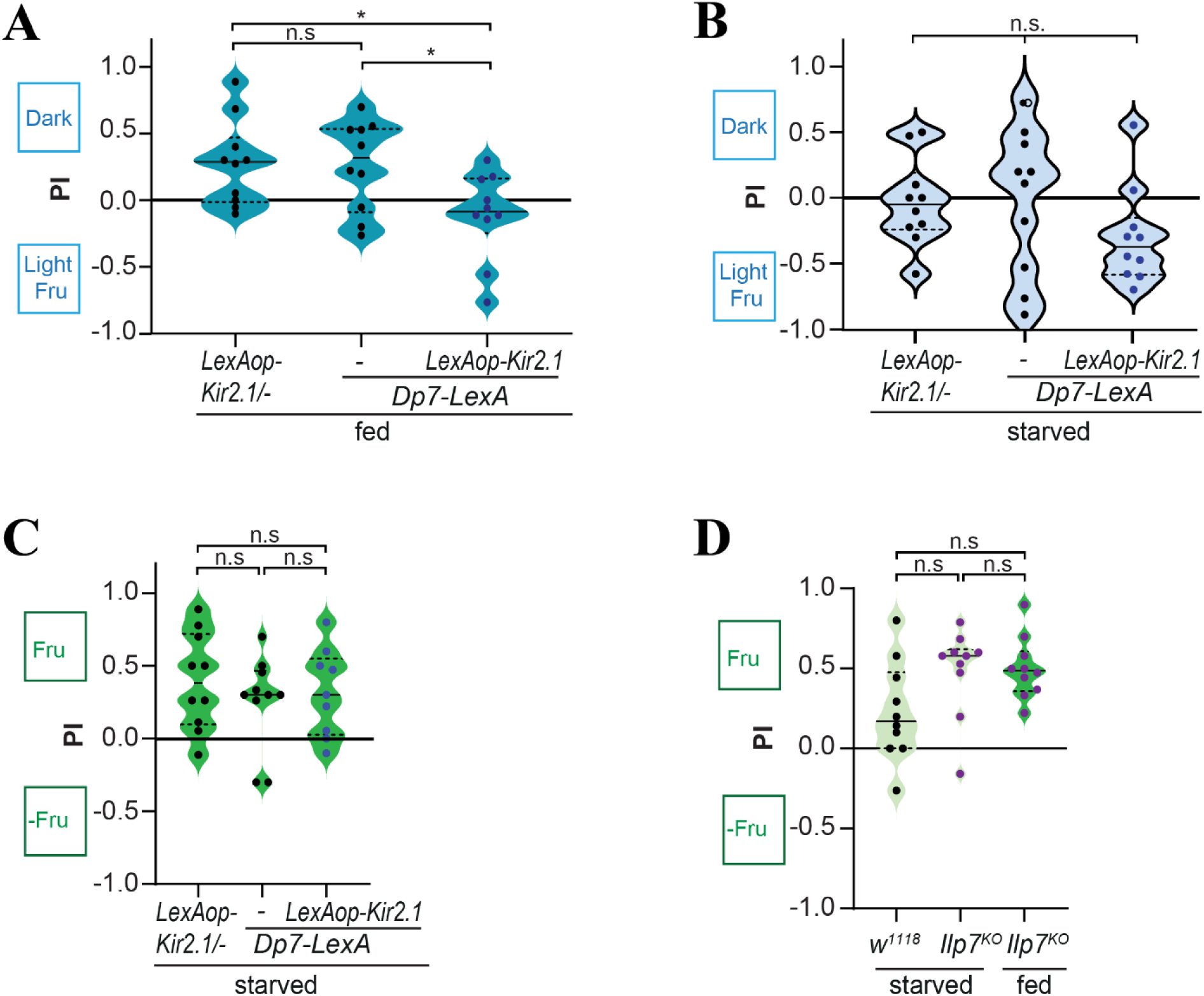
**A.** Silencing of Dp7 neurons (*Dp7-LexA*>*LexAop-Kir2.1*) in fed animals increased preference towards light and fructose compared to controls (n=10 trials/genotype, one-way ANOVA with Tukey’s post hoc test, *P < 0.05, n.s, non-significant). **B.** Silencing of Dp7 neurons (*Dp7-LexA*>*LexAop-Kir2.1*) in starved animals showed a similar preference to light and fructose as the controls. (n=10 trials/genotype, one-way ANOVA with Tukey’s post-hoc test, *P < 0.05, n.s, non-significant). **C.** Silencing of Dp7 neurons (*Dp7-LexA*>*LexAop-Kir2.1*) in starved animals showed similar fructose preference behavior as controls. (n=10 trials/genotype, one-way ANOVA with Tukey’s post-hoc test, n.s, non-significant). **D.** *Ilp7^ko^* larvae showed a similar preference for fructose as w*^1118^* animals, independent of the feeding state (n=10 trials/genotype one-way ANOVA with Tukey’s post-hoc test, n.s, non-significant).

**Extended Fig. 4.**
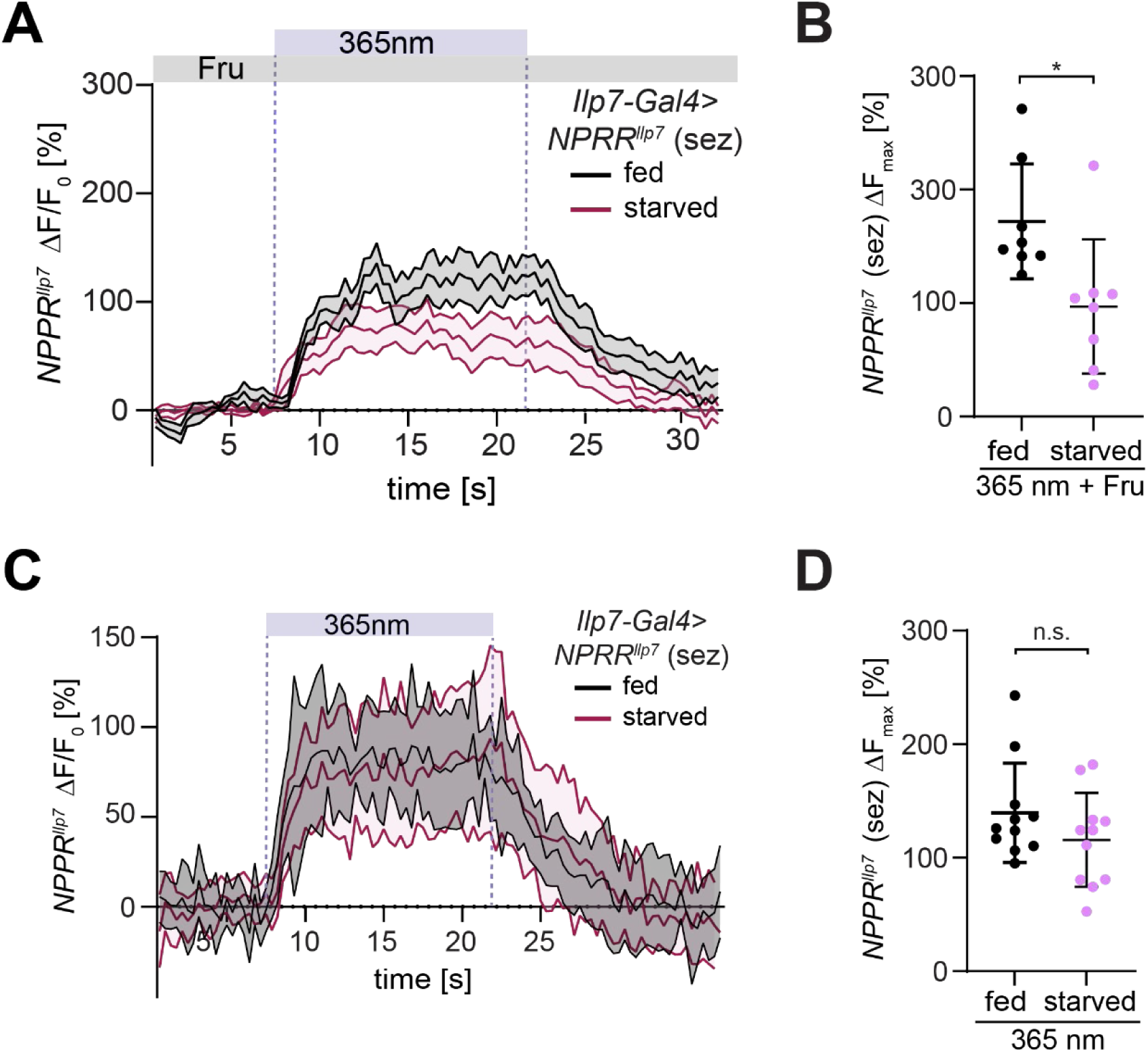
**A.** Average fluorescent change of NPRR^Ilp7^ signals in Dp7 neurons in the sez region (*Ilp7-Gal4>NPRR^ilp7^*) in response to light and in the presence of fructose (0.1M). Fed and starved animals differ in their capacity for Ilp7 release. Dotted lines show the stimulation period with light (mean±SEM, n=8,7 neuropeptide puncta from 4 animals per condition). **B.** Maximum fluorescence changes (ΔF_max_) of NPRR^Ilp7^ signals in response to light and fructose in fed and starved larvae (n=8,7 neuropeptide puncta from 4 animals per condition, unpaired t-test with Welch’s correction, **P<0.01). **C.** Average fluorescent change of NPRR^Ilp7^ signals in Dp7 neurons in the sez region (*Ilp7-Gal4>NPRR^ilp7^*) in response to light without fructose. Fed and starved animals show similar capacities for Ilp7 release without fructose. Dotted lines show the stimulation period with light (mean±SEM, n=11,11 neuropeptide puncta from 3 animals per condition). **D.** Maximum fluorescence changes (ΔF_max_) of NPRR^Ilp7^ signals in response to light without fructose in fed and starved larvae (n=11, 11 neuropeptide puncta from 3 animals per condition, unpaired t-test with Welch’s correction, n.s. P>0.05).

**Extended Figure 6.**
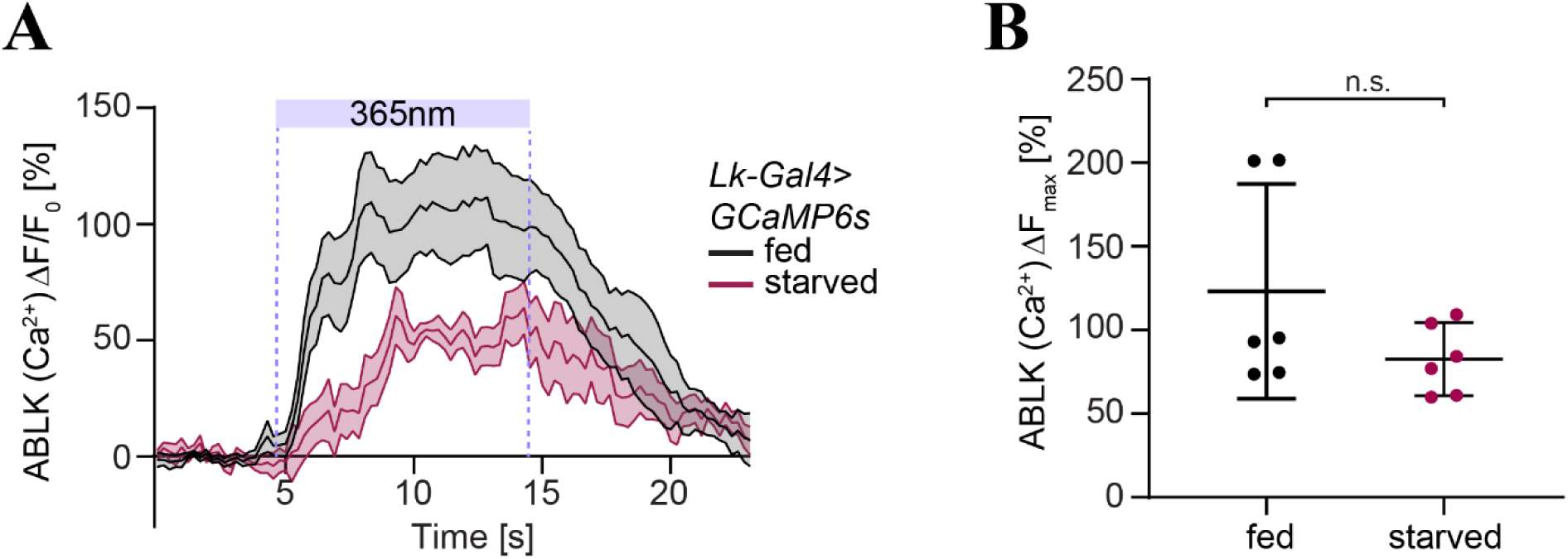
**A.** Average fluorescent change of ABLK neuron calcium (GCaMP6s) signals in fed and starved larvae in response to 365 nm light (*Lk-Gal4>UAS-GCaMP6s,* mean±SEM, n=6 animals). **B.** Maximum fluorescence changes (ΔF_max_) of ABLK calcium signals from **A** (n=5 animals, unpaired t-test with Welch’s correction, n.s., non-significant).

## Notes

### Competing Interest Statement

The authors have declared no competing interest.

## References

1. Sugrue, L. P., Corrado, G. S. & Newsome, W. T. Choosing the greater of two goods: Neural currencies for valuation and decision making. Nat Rev Neurosci 6, 363–375 (2005).

2. Stein, B. E. & Stanford, T. R. Multisensory integration: Current issues from the perspective of the single neuron. Nat Rev Neurosci 9, 255–266 (2008).

3. Thapliyal, S., Beets, I. & Glauser, D. A. Multisite regulation integrates multimodal context in sensory circuits to control persistent behavioral states in C. elegans. Nature Communications 2023 14:1 14, 1–19 (2023).

4. Flavell, S. W., Gogolla, N., Lovett-Barron, M. & Zelikowsky, M. The emergence and influence of internal states. Neuron 110, 2545–2570 (2022).

5. Grunwald Kadow, I. C. State-dependent plasticity of innate behavior in fruit flies. Curr Opin Neurobiol 54, 60–65 (2019).

6. Smith, N. K. & Grueter, B. A. Hunger-driven adaptive prioritization of behavior. FEBS J 289, 922–936 (2022).

7. Tinbergen, N. The Study of Instinct. The study of instinct. (Clarendon Press/Oxford University Press, New York, NY, US, 1951).

8. Münch, D., Ezra-Nevo, G., Francisco, A. P., Tastekin, I. & Ribeiro, C. Nutrient homeostasis — translating internal states to behavior. Curr Opin Neurobiol 60, 67–75 (2020).

9. Andermann, M. L. & Lowell, B. B. Toward a Wiring Diagram Understanding of Appetite Control. Neuron 95, 757–778 (2017).

10. Miroschnikow, A., Schlegel, P. & Pankratz, M. J. Making Feeding Decisions in the Drosophila Nervous System. Current Biology 30, R831–R840 (2020).

11. Kim, S. K., Tsao, D. D., Suh, G. S. B. & Miguel-Aliaga, I. Discovering signaling mechanisms governing metabolism and metabolic diseases with Drosophila. Cell Metab 33, 1279–1292 (2021).

12. Nässel, D. R. & Zandawala, M. Endocrine cybernetics: neuropeptides as molecular switches in behavioural decisions. Open Biol 12, 24–26 (2022).

13. Nässel, D. R. & Zandawala, M. Hormonal axes in Drosophila: regulation of hormone release and multiplicity of actions. Cell Tissue Res 382, 233–266 (2020).

14. Dus, M. et al. Nutrient Sensor in the Brain Directs the Action of the Brain-Gut Axis in Drosophila. Neuron 87, 139–151 (2015).

15. Yang, Z. et al. A post-ingestive amino acid sensor promotes food consumption in Drosophila. Cell Research 2018 28:10 28, 1013–1025 (2018).

16. Poe, A. R. et al. Developmental emergence of sleep rhythms enables long-term memory in Drosophila. Sci Adv 9, (2023).

17. Cavanaugh, D. J. et al. Identification of a Circadian Output Circuit for Rest:Activity Rhythms in Drosophila. Cell 157, 689–701 (2014).

18. Filosa, A., Barker, A. J., Dal Maschio, M. & Baier, H. Feeding State Modulates Behavioral Choice and Processing of Prey Stimuli in the Zebrafish Tectum. Neuron 90, 596–608 (2016).

19. Alhadeff, A. L. et al. A Neural Circuit for the Suppression of Pain by a Competing Need State. Cell 173, 140–152.e15 (2018).

20. Oh, Y. & Suh, G. S. B. Starvation-induced sleep suppression requires the Drosophila brain nutrient sensor. J Neurogenet 37, 70–77 (2023).

21. Lewis, L. P. C. et al. A Higher Brain Circuit for Immediate Integration of Conflicting Sensory Information in Drosophila. Current Biology 25, 2203–2214 (2015).

22. Bräcker, L. B. et al. Essential role of the mushroom body in context-dependent CO_2_ avoidance in drosophila. Current Biology 23, 1228–1234 (2013).

23. LeDue, E. E. et al. Starvation-Induced Depotentiation of Bitter Taste in Drosophila. Current Biology (2016) doi:10.1016/j.cub.2016.08.028.

24. Sareen, P. F., McCurdy, L. Y. & Nitabach, M. N. A neuronal ensemble encoding adaptive choice during sensory conflict in Drosophila. Nature Communications 2021 12:1 12, 1–13 (2021).

25. Lin, H. et al. A nutrient-specific gut hormone arbitrates between courtship and feeding. Nature 602, 632–638 (2022).

26. Mishra, D. et al. The molecular basis of sugar sensing in drosophila larvae. Current Biology 23, 1466–1471 (2013).

27. Imambocus, B. N. et al. A neuropeptidergic circuit gates selective escape behavior of Drosophila larvae. Current Biology 32, 149–163.e8 (2022).

28. Humberg, T.-H. & Sprecher, S. G. Age- and Wavelength-Dependency of Drosophila Larval Phototaxis and Behavioral Responses to Natural Lighting Conditions. Front Behav Neurosci 11, 1–13 (2017).

29. Keene, A. C. et al. Distinct Visual Pathways Mediate Drosophila Larval Light Avoidance and Circadian Clock Entrainment. Journal of Neuroscience 31, 6527–6534 (2011).

30. Busto, M., Iyengar, B. & Campos, a R. Genetic dissection of behavior: modulation of locomotion by light in the Drosophila melanogaster larva requires genetically distinct visual system functions. J Neurosci 19, 3337–44 (1999).

31. Fujita, M. & Tanimura, T. Drosophila Evaluates and Learns the Nutritional Value of Sugars. Current Biology 21, 751–755 (2011).

32. Rohwedder, A. et al. Nutritional value-dependent and nutritional value-independent effects on Drosophila melanogaster larval behavior. Chem Senses 37, 711–721 (2012).

33. Oh, Y. & Suh, G. S. B. Starvation-induced sleep suppression requires the Drosophila brain nutrient sensor. J Neurogenet 37, 70–77 (2023).

34. Hückesfeld, S. et al. Unveiling the sensory and interneuronal pathways of the neuroendocrine connectome in Drosophila. Elife 10, 2020.10.22.350306 (2021).

35. Masuyama, K., Zhang, Y., Rao, Y. & Wang, J. W. Mapping Neural Circuits with Activity-Dependent Nuclear Import of a Transcription Factor. J Neurogenet 26, 89–102 (2012).

36. Xiang, Y. et al. Light-avoidance-mediating photoreceptors tile the Drosophila larval body wall. Nature 468, 921–6 (2010).

37. Hu, C. et al. Sensory integration and neuromodulatory feedback facilitate Drosophila mechanonociceptive behavior. Nat Neurosci 20, 1085–1095 (2017).

38. Schlegel, P. et al. Synaptic transmission parallels neuromodulation in a central food-intake circuit. Elife 5, 462–465 (2016).

39. Li, K. et al. Belly roll, a GPI-anchored Ly6 protein, regulates Drosophila melanogaster escape behaviors by modulating the excitability of nociceptive peptidergic interneurons. Elife 12, (2023).

40. Inagaki, H. K., Panse, K. M. & Anderson, D. J. Independent, reciprocal neuromodulatory control of sweet and bitter taste sensitivity during starvation in Drosophila. Neuron 84, 806–820 (2014).

41. Vogt, K. et al. Internal state configures olfactory behavior and early sensory processing in Drosophila larvae. Sci Adv 7, eabd6900 (2021).

42. Devineni, A., Sun, B., Zhukovskaya, A. & Axel, R. Acetic acid activates distinct taste pathways in Drosophila to elicit opposing, state-dependent feeding responses. 1–23 (2018) 10.1101/378414.

43. Humberg, T. H. et al. Dedicated photoreceptor pathways in Drosophila larvae mediate navigation by processing either spatial or temporal cues. Nature Communications 2018 9:1 9, 1–16 (2018).

44. Gomez-Marin, A., Stephens, G. J. & Louis, M. Active sampling and decision making in Drosophila chemotaxis. Nat. Commun. 2, (2011).

45. Kane, E. A., et al. Sensorimotor structure of Drosophila larva phototaxis. Proc. Natl Acad. Sci. USA 110, E3868–E3877 (2013).

46. Paisios, E., Rjosk, A., Pamir, E. & Schleyer, M. Common microbehavioral ‘footprint’ of two distinct classes of conditioned aversion. Learning and Memory 24, 191–198 (2017).

47. Schleyer, M. et al. The impact of odor–reward memory on chemotaxis in larval Drosophila. Learning & Memory 22, 267 (2015).

48. Du, E. J. et al. Nucleophile sensitivity of Drosophila TRPA1 underlies light-induced feeding deterrence. Elife 5, (2016).

49. Guntur, A. R., et al. *Drosophila* TRPA1 isoforms detect UV light via photochemical production of H _2_ O _2_. Proceedings of the National Academy of Sciences 201514862 (2015) doi:10.1073/pnas.1514862112.

50. Cheriyamkunnel, S. J. et al. A neuronal mechanism controlling the choice between feeding and sexual behaviors in Drosophila. Current Biology 31, 4231–4245.e4 (2021).

51. Cazalé-Debat, L. et al. Mating proximity blinds threat perception. Nature 634, (2024).

52. Jovanoski, K. D. et al. Dopaminergic systems create reward seeking despite adverse consequences. Nature 2023 623:7986 623, 356–365 (2023).

53. Held, M. et al. Aminergic and peptidergic modulation of Insulin-Producing Cells in Drosophila. Elife 13, (2024).

54. Bisen, R. S., Iqbal, F. M., Cascino-Milani, F., Bockemühl, T. & Ache, J. M. Nutritional state-dependent modulation of Insulin-Producing Cells in Drosophila. Elife 13, (2024).

55. Oh, Y. et al. Periphery signals generated by Piezo-mediated stomach stretch and Neuromedin-mediated glucose load regulate the Drosophila brain nutrient sensor. Neuron 109, 1979–1995.e6 (2021).

56. Betley, J. N., Cao, Z. F. H., Ritola, K. D. & Sternson, S. M. Parallel, Redundant Circuit Organization for Homeostatic Control of Feeding Behavior. Cell 155, 1337–1350 (2013).

57. Burnett, C. J. et al. Hunger-Driven Motivational State Competition. Neuron 92, 187–201 (2016).

58. Padilla, S. L. et al. Agouti-related peptide neural circuits mediate adaptive behaviors in the starved state. Nat Neurosci 19, 734–741 (2016).

59. Comeras, L. B., Herzog, H. & Tasan, R. O. Neuropeptides at the crossroad of fear and hunger: a special focus on neuropeptide Y. Ann N Y Acad Sci 1455, 59–80 (2019).

60. Tseng, Y. T., Schaefke, B., Wei, P. & Wang, L. Defensive responses: behaviour, the brain and the body. Nature Reviews Neuroscience 2023 24:11 24, 655–671 (2023).

61. Zandawala, M., Marley, R., Davies, S. A. & Nässel, D. R. Characterization of a set of abdominal neuroendocrine cells that regulate stress physiology using colocalized diuretic peptides in Drosophila. Cellular and Molecular Life Sciences 75, 1099–1115 (2018).

62. Cannell, E. et al. The corticotropin-releasing factor-like diuretic hormone 44 (DH44) and kinin neuropeptides modulate desiccation and starvation tolerance in Drosophila melanogaster. Peptides (N.Y.) 80, 96–107 (2016).

63. Johnson, E. C. et al. A novel diuretic hormone receptor in Drosophila: evidence for conservation of CGRP signaling. Journal of Experimental Biology 208, 1239–1246 (2005).

64. Chatzigeorgiou, M. & Schafer, W. R. Lateral Facilitation between Primary Mechanosensory Neurons Controls Nose Touch Perception in C. elegans. Neuron 70, 299–309 (2011).

65. Gambino, F. et al. Sensory-evoked LTP driven by dendritic plateau potentials in vivo. Nature 515, 116–119 (2014).

66. Chen, Y., Lin, Y. C., Kuo, T. W. & Knight, Z. A. Sensory Detection of Food Rapidly Modulates Arcuate Feeding Circuits. Cell 160, 829–841 (2015).

67. Hu, Y. et al. A Neural Basis for Categorizing Sensory Stimuli to Enhance Decision Accuracy. Current Biology 30, 4896–4909.e6 (2020).

68. Katzen, A. et al. The nematode worm C. Elegans chooses between bacterial foods as if maximizing economic utility. Elife 12, (2023).

69. Deng, B. et al. Chemoconnectomics: Mapping Chemical Transmission in Drosophila. Neuron 101, 876–893.e4 (2019).

70. Yang, C.-H., Belawat, P., Hafen, E., Jan, L. Y. & Jan, Y.-N. Drosophila egg-laying site selection as a system to study simple decision-making processes. Science 319, 1679–83 (2008).

71. Masuyama, K., Zhang, Y., Rao, Y. & Wang, J. W. Mapping Neural Circuits with Activity-Dependent Nuclear Import of a Transcription Factor. J Neurogenet 26, 89–102 (2012).

72. Grönke, S., Clarke, D.-F., Broughton, S., Andrews, T. D. & Partridge, L. Molecular evolution and functional characterization of Drosophila insulin-like peptides. PLoS Genet 6, e1000857 (2010).

73. Baines, R. A., Uhler, J. P., Thompson, A., Sweeney, S. T. & Bate, M. Altered electrical properties in Drosophila neurons developing without synaptic transmission. J Neurosci 21, 1523–1531 (2001).

74. de Haro, M. et al. Detailed analysis of leucokinin-expressing neurons and their candidate functions in the Drosophila nervous system. Cell Tissue Res 339, 321–336 (2010).

75. Simpson, J. H. Rationally subdividing the fly nervous system with versatile expression reagents. J Neurogenet 30, 185–194 (2016).

76. Imambocus, B. N., Formozov, A., Zhou, F. & Soba, P. A two-choice assay for noxious light avoidance with temporal distribution analysis in Drosophila melanogaster larvae. STAR Protoc 3, 101787 (2022).

77. Thane, M. et al. High-resolution analysis of individual Drosophila melanogaster larvae uncovers individual variability in locomotion and its neurogenetic modulation. Open Biol 13, (2023).

78. Saalfeld, S., Cardona, A., Hartenstein, V. & Tomancak, P. CATMAID: collaborative annotation toolkit for massive amounts of image data. Bioinformatics 25, 1984–1986 (2009).

